# Ga_q_-mediated contact release is required for efficient mitochondrial quality control

**DOI:** 10.64898/2026.10.30.761943

**Authors:** Miriam Ferrando-Soler, Georgina Garrido, Mireia Andreu-Carbó, Alex Ascunce-París, Javier Rubio, Alba Benages, Ismael Izquierdo-Villalba, Federico Mayor, Catalina Ribas, Carlos Enrich, Carles Rentero, Cristiane Benincá, Anna M. Aragay

## Abstract

Lysosomes and mitochondria are highly dynamic organelles whose coordinated interactions are essential for cellular homeostasis, while their dysfunction contributes to numerous human diseases. Here, we identify the mitochondrial outer membrane protein Fis1 and the RabGAPs TBC1D15 and TBC1D17 as novel interacting partners of the heterotrimeric G protein a-subunit Ga_q_. Using mitochondrial proteomics, co-immunoprecipitation, proximity ligation assays, and in vitro binding studies with purified proteins, we demonstrate that Ga_q_ forms a complex with Fis1 and TBC1D15/TBC1D17 through a non-canonical interaction interface. Ga_q_ localizes to mitochondria-lysosome contact sites, where it regulates the dynamic balance between contact formation and dissociation. Loss of Ga_q_ resulted in enlarged and clustered lysosomes and increased the extent and duration of mitochondria-lysosome contacts. Live-cell imaging revealed prolonged organelle tethering in Ga_q_-deficient cells, while proximity ligation and three-dimensional reconstruction analyses localized Ga_q_ and TBC1D15 to mitochondrial tips closely associated with lysosomes. This localization, together with increased mitochondrial fission and altered mitophagic turnover following Ga_q_ loss, links the regulation of contacts at mitochondrial tips to mitochondrial fragmentation and quality control. Re-expression of Ga_q_, but not the non-canonical interaction-deficient mutant, restored normal lysosomal morphology, contact-site dynamics, mitochondrial fragmentation, and mitophagy. Together, these findings identify a Ga_q_ as a dynamic regulator of mitochondria-lysosome untethering and connect contact site turnover with mitochondria fragmentation and mitophagy. Our work uncovers a novel link between G protein signaling, organelle communication, and mitochondrial quality control.

## Introduction

Recently, numerous GPCRs and their associated G proteins have been identified at cellular endomembranes, expanding their functional significance beyond the plasma membrane (Crilly & Puthenveedu, 2021; Eichel & von Zastrow, 2018; Fasciani *et al*, 2022; Hewavitharana & Wedegaertner, 2012; Jong *et al*, 2018; O’Malley *et al*, 2003; Radoux-Mergault *et al*, 2023; Zhang *et al*, 2012). Multiple studies have now demonstrated that mitochondrial GPCRs (mito-GPCRs) influence oxidative activity, reactive oxygen species (ROS) production, respiration, apoptosis, nitric oxide synthesis, and calcium transport (Fasciani *et al*, 2022; Hewavitharana & Wedegaertner, 2012; Mendizabal-Zubiaga *et al*, 2016; Picard & Shirihai, 2022). Ga_q_ is a well-characterized heterotrimeric G protein, primarily known for its canonical role in membrane receptor (GPCR) signaling through phosphoinositide phospholipase C□ and its involvement in a wide range of physiological processes and cancers (Kamato *et al*, 2017; Litosch, 2016; Navarro-Lérida *et al*, 2022). Ga_q_ has also been located in endomembranes, particularly in mitochondria and lysosomes, where it has been found to play a pivotal role in regulating cellular physiology (Benincá *et al*, 2014; Cabezudo *et al*, 2021). In mitochondria, Ga_q_ influences fusion and fission processes, mitochondrial morphology, respiratory chain activity, and the assembly of supercomplexes (Benincá *et al*, 2014; Izquierdo-Villalba *et al*, 2024), underscoring its critical role in mitochondrial function and cellular metabolism. However, distinguishing Ga_q_’s functions at endomembranes from its signaling from GPCRs at the plasma membrane has been challenging. The finding that mitochondrial proteins Rho-like Miro1/2 and the armadillo protein ArmcX3 (Alex3) interacts with Ga_q_ has help to define its role in mitochondria (Izquierdo-Villalba *et al*, 2024). This interaction is essential for regulating mitochondrial movement along microtubules in hippocampal neuron axons. Moreover, Ga_q_ subunit has been demonstrated to regulate autophagy by promoting mTOR signaling (Cabezudo *et al*, 2021). However, it remains to be stablished how Ga_q_ signaling governs the morphology, distribution, and interplay between mitochondria and other organelles and the impact in cell physiology.

Notably, membrane contact sites are increasingly recognized as essential for inter-organelle communication and regulation. Their importance in maintaining lysosome distribution and function, mitophagy, and overall cellular homeostasis is well established (Wong *et al*, 2018, 2019, 2022). Moreover, recent studies have begun to highlight the role of these contacts in several diseases, such as Charcot-Marie-Tooth disease, Parkinson’s disease, Alzheimer’s disease, and lysosomal storage disorders (Cisneros *et al*, 2022; Girolimetti *et al*, 2025; Kim *et al*, 2021; Wong *et al*, 2019). The contacts between mitochondria and lysosomes are finely regulated by tethering and untethering protein machinery, primarily controlled by the Rab7 GTPase (Wong *et al*, 2018). The recruitment of the RabGAP protein TBC1D15 by the mitochondrial outer membrane protein Fis1 promotes the inactivation of Rab7 on membranes, which, in turn, facilitates the detachment of lysosomes from mitochondria (Wong *et al*, 2018). Importantly, mitochondrial fission can support distinct cellular outcomes depending on the position of division. Whereas midzone fission is associated with mitochondrial proliferation, peripheral fission can segregate damaged mitochondrial material into smaller daughter mitochondria for subsequent removal by mitophagy. Peripheral fission is preceded by lysosomal contact and is regulated by Fis1, providing a functional connection between mitochondria-lysosome interactions, mitochondrial fragmentation, and quality control (Kleele *et al*, 2021). These findings raise the question of how the regulation of contact-site duration and dissociation is coordinated with mitochondrial fragmentation and mitophagic turnover.

Here, we identify a previously unrecognized role for Ga_q_ in the regulation of mitochondria-lysosome communication. In a proteomic analysis combined with molecular assays, we have identified the mitochondrial protein Fis1 and the proteins TBC1D15 and TBC1D17 as interactor proteins for Ga_q_. Proximity ligation assays (PLA) localized Ga_q_ at mitochondria tips, together with TBC1D15, and Lys-LE, highlighting its implication in the regulation of mitochondrial fragmentation and mitophagy. Loss of Ga_q_ results in sustained mitochondria-lysosome associations, lysosomal clustering, mitochondrial fragmentation, and enhanced mitophagic flux, whereas disruption of the Ga_q_-TBC1D15/Fis1 interaction phenocopies these defects. Collectively, our findings uncover a non-canonical function of Ga_q_ as a key organizer of interorganelle communication and establish the Ga_q_-Fis1-TBC1D15 axis as a critical regulatory node linking mitochondria-lysosome contacts to mitochondrial quality control.

## Results

### Ga_q_ interacts with mitochondrial protein Fis1 and TBC1D15/TBC1D17

To identify novel regulators of mitochondrial dynamics and mitophagy associated with Ga_q_ signaling, we revisited a previously generated endomembrane/mitochondrial-enriched proteomic dataset (Izquierdo-Villalba *et al*, 2024). This analysis compared membrane fractions from wild-type murine embryonic fibroblasts (WT), Ga_q/11_-deficient MEFs (Ga_q/11_ KO), Ga_q_-reconstituted KO MEFs (KI), and NIH3T3 cells subjected to immunoprecipitation with two independent anti-Ga_q_ antibodies followed by mass spectrometry analysis. Among the proteins selectively detected in Ga_q_-containing samples but absent in Ga_q/11_ KO cells were several factors involved in mitochondrial quality control, mitophagy, and lysosomal regulation (Izquierdo-Villalba *et al*, 2024). Notably, the proteins Fis1 and TBC1D17 were identified in WT MEFs and NIH3T3 cells but were absent from Ga_q/11_ KO samples.

We validated the proteomic result by immunoprecipitating endogenous Ga_q_ with Fis1 from MEF and HEK293 cells (Fig. 1A and B). In addition, HA-tagged Fis1 efficiently co-immunoprecipitated with Ga_q_ (Fig. 1C). Because Fis1 is known to interact with TBC1D15 and TBC1D17, both key regulators of mitochondria-lysosome contacts and mitophagy (Wong *et al*, 2018; Yamano *et al*, 2014), we investigated whether Ga_q_ also associates with these proteins. Moreover, as mentioned, TBC1D17 is also present in the Gq-mitoproteomic analysis (Izquierdo-Villalba *et al*, 2024). TBC1D15 was detected in HA-Fis1 immunocomplexes in the presence of active Ga_q_ (Ga_q_R183C) (Fig. S1A), suggesting the existence of a common protein assembly. Constitutive active Ga_q_ immunoprecipitated with endogenous TBC1D15 from cellular lysates (Fig. 1D). Consistent with this observation, both Ga_q_ and endogenous TBC1D15 were detected in Flag-TBC1D17 immunoprecipitates (Fig. 1E), while Flag-tagged TBC1D17 efficiently co-immunoprecipitated with antibodies against Ga_q_ (Fig. S1B). In contrast, no interaction was detected between TBC1D15 and the mitochondrial Gβγ dimer (Gβ_1_γ_2_) demonstrating the specificity of the association with Ga_q_ and indicating that TBC1D15 preferentially associates with the Ga_q_ subunit rather than with heterotrimeric G□□□ components (Fig. S1C). Collectively, these findings suggest that Ga_q_ may be a novel binding partner of Fis1 and its associated TBC1D15 and TBC1D17 RabGAPs.

**Figure 1:**
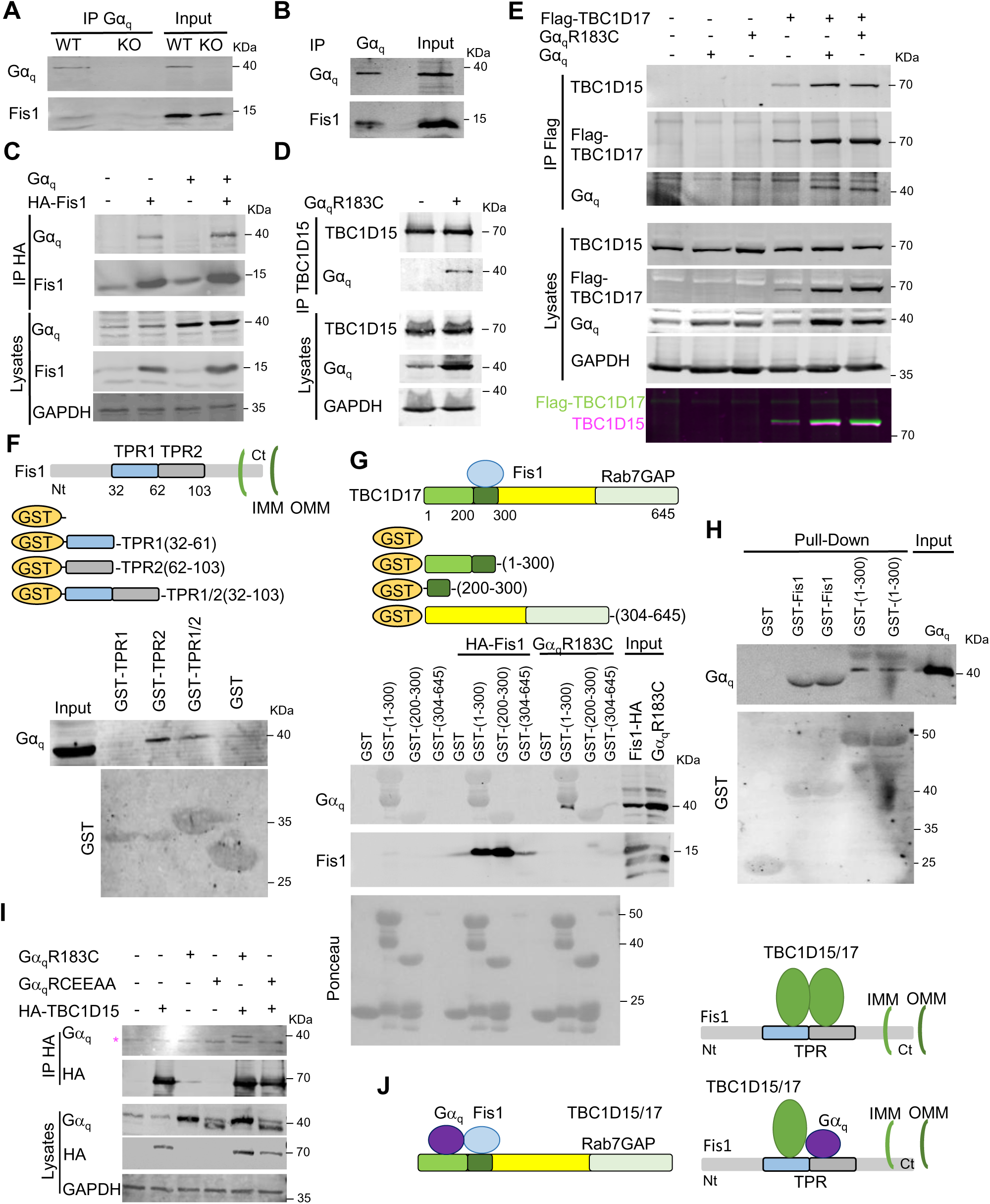
Ga_q_ interacts with Fis1, TBC1D15 and TBC1D17. (**A**) WT and *Ga_q/11_*^-/-^ (KO) MEF homogenates were immunoprecipitated with anti-Ga_q_ and immunoblotted for Ga_q_ and Fis1. Input is shown. (**B**) Endogenous Ga_q_ immunoprecipitates from HEK293 cells immunoblotted for Ga_q_ and Fis1. Input is shown. (**C**) HA-immunoprecipitates and lysates from HEK293 cells expressing HA-Fis1 and Ga_q_, immunoblotted for Fis1, Ga_q_ and GAPDH. (**D**) Endogenous TBC1D15 immunoprecipitates and lysates from HEK293 cells expressing Ga_q_R183C, immunoblotted for TBC1D15, Ga_q_ and GAPDH. (**E**) Flag-immunoprecipitates and lysates from HEK293 cells expressing Flag-TBC1D17, Ga_q_ or Ga_q_R183C, immunoblotted for Flag, TBC1D15, Ga_q_ and GAPDH. (**F**) (Schematic representation of Fis1 and GST-fused Fis1 peptides. Pull-downs using GST, GST-TPR1 (aa 32-61), GST-TPR2 (aa 62-103) or GST-TPR1/2 (aa 32-103) from HEK293 lysates, immunoblotted for Ga_q_. Ponceau staining and input are shown. (**G**) Schematic representation of TBC1D17 domain organization and GST-fused peptides. Pull-downs using GST-TBC1D17 fragments (aa 1-300, 200-300 or 304-645) from HEK293 lysates expressing HA-Fis1 and Ga_q_R183C, immunoblotted for Ga_q_ and Fis1. Anti-GST staining is shown. (**H**) Pull-downs of purified Ga_q_ with GST, GST-Fis1 or GST-TBC1D17 (fragment aa 1-300), immunoblotted for Ga_q_ and GST. Input shows purified Ga_q_. (**I**) HA-immunoprecipitates and lysates from HEK293 cells expressing HA-TBC1D15, Ga_q_R183C or Ga_q_R183C/E234A/E245A (Ga_q_RCEEAA), immunoblotted for HA, Ga_q_ and GAPDH. The magenta asterisk in the IP bands indicates a nonspecific band. (**J**) Model of Fis1 interaction with TBC1D15 and/or TBC1D17 at the mitochondrial outer membrane in the absence or presence of Ga_q_. All representative blots from ≥3 independent experiments.

To gain mechanistic insight into complex assembly, we mapped the interaction domains involved. Fis1, highly conserved from yeast to mammals, anchors to the outer mitochondrial membrane through its C-terminal transmembrane region (Mozdy *et al*, 2000). It interacts with TBC1D15 and TBC1D17 through its cytosolic N-terminal tetratricopeptide repeat (TPR) domain (Suzuki *et al*, 2003). Ga_q_ was found to bind a proximal region within the Fis1 TPR domain encompassing amino acids 62-103 (TPR2) (Fig. 1F). Notably, this region lies within the broader Fis1 segment previously reported to mediate TBC1D15/TBC1D17 binding and oligomerization (Onoue *et al*, 2013; Yamano *et al*, 2014). Conversely, Fis1 interacted with residues 200-300 of TBC1D17, corresponding to the homologous Fis1-binding region previously described for TBC1D15. In contrast, Ga_q_ associated with the adjacent N-terminal region of TBC1D17 (amino acids 1-200) (Fig. 1G, S1D). Interestingly, this N-terminal region in TBC1D15 and TBC1D17 contains a PB1-like domain (amino acids 42-196), besides an unstructured loop (amino acids 56-122) predicted by AlphaFoold (Fig. S1D). These mapping studies suggest that Ga_q_, Fis1, and TBC1D15/TBC1D17 occupy neighboring but non-overlapping binding surfaces, supporting the formation of a heterotrimeric protein complex. To determine whether these interactions were direct, purified recombinant Ga_q_ protein (Chan *et al*, 2011) was incubated with GST-fused forms of Fis1 and TBC1D17-binding region (1-300). Pull-down experiments demonstrated direct binding of Ga_q_ to both Fis1 and TBC1D17 (Fig. 1H).

Previous work identified a non-canonical PB1-like domain within Ga_q_ that participates in PB1-type of protein interactions and in the regulation of autophagy independently of classical G-protein effector signaling (Cabezudo *et al*, 2021). To determine whether this region mediates interaction with TBC proteins, we examined a PB1-like domain mutant in the Ga_q_ constitutive active form (Ga_q_R183C/E234A/E245A; also named Ga_q_RCEEAA). This mutant exhibited markedly reduced co-immunoprecipitation with TBC1D15 compared with wild-type Ga_q_, indicating that TBC1D15 recognition requires the integrity of this non-canonical interaction surface (Fig. 1I).

Taken together, these results support a model in which Ga_q_ forms a macromolecular complex with Fis1 and TBC1D15 and TBC1D17 at the mitochondrial outer membrane (Fig. 1J).

### Ga_q_ Localizes in Close Proximity to TBC1D15 at Endolysosomal Compartments

To investigate the subcellular localization of Ga_q_ and TBC1D15, we first examined the distribution of the endogenous proteins in COS7 cells. TBC1D15 was detected throughout the cytoplasm in a punctate patter, with pronounced enrichment in perinuclear regions (Fig. S2A). Similarly, Ga_q_ localized along cytoplasmic and accumulated in the perinuclear area (Fig. S2A). In selected regions, both proteins were found in close apposition at discrete punctate sites both in endogenous and overexpressed Ga_q_ (Fig. S2A) presenting some co-localiztion (Fig. S2B). Expression of HA-tagged TBC1D15 recapitulated the localization pattern of the endogenous protein and further highlighted its close spatial association with Ga_q_ (Fig. S2C). Deconvolution microscopy revealed highly organized subcellular domains containing punctate structures that follows a linear structure in which Ga_q_ and TBC1D15 were separated by only short distances (Fig. S2C). Consistently, line-scan analyses confirmed the close juxtaposition of both proteins at these sites (Fig. S2D).

To further characterize the subcellular context of these interactions, late endosome-lysosome (LE-Lys) compartments were labeled with anti-LAMP1 antibodies (Fig. 2A-C). High-resolution imaging revealed that TBC1D15 formed discrete clusters aligned along the membrane of LAMP1-positive vesicles where Ga_q_ was also present (Fig.2A, B). Further analysis using 3D reconstruction revealed the close apposition between TBC1D15 and Ga_q_ within LE-Lys compartments (Fig. 2 C). To directly assess the proximity between these proteins, we performed Proximity Ligation Assays (PLA). The analysis using antibodies against HA-tagged TBC1D15 and endogenous Ga_q_ generated robust signals that were detected only when both primary antibodies were close (less than 40 nm) and with both antibodies present in the samples (Fig. 2D, Fig. 2SE). Moreover, these signals were absent in untransfected (without HA-TBC1D15 expression) MEF cells and in absence of Ga_q/11_ in Ga_q/11_KO MEF cells, confirming assay specificity (Fig. 2D). The assay was also validated in HEK293 expressing HA-TBC1D15 and endogenous Ga_q_ (Fig.2SF). Similarly, PLA performed between HA-tagged Fis1 and endogenous Ga_q_ revealed specific interaction signals, although fewer puncta were observed compared with the TBC1D15-Ga_q_ pair (Fig. 2E), consistent with a reduced mitochondrial localization of Ga_q_ (Benincá *et al*, 2014). Appropriate negative controls confirmed the specificity of these interactions (Fig. 2E). To further confirm these results, PLA samples (Ga_q_-TBC1D15 interactions) were further treated with antibodies against HA to visualize TBC1D15 location in the cell. PLA puncta corresponding to Ga_q_-TBC1D15 interactions (in magenta) were detected in HA-TBC1D15 labelled signals (green) (Fig. 2F).

**Figure 2.**
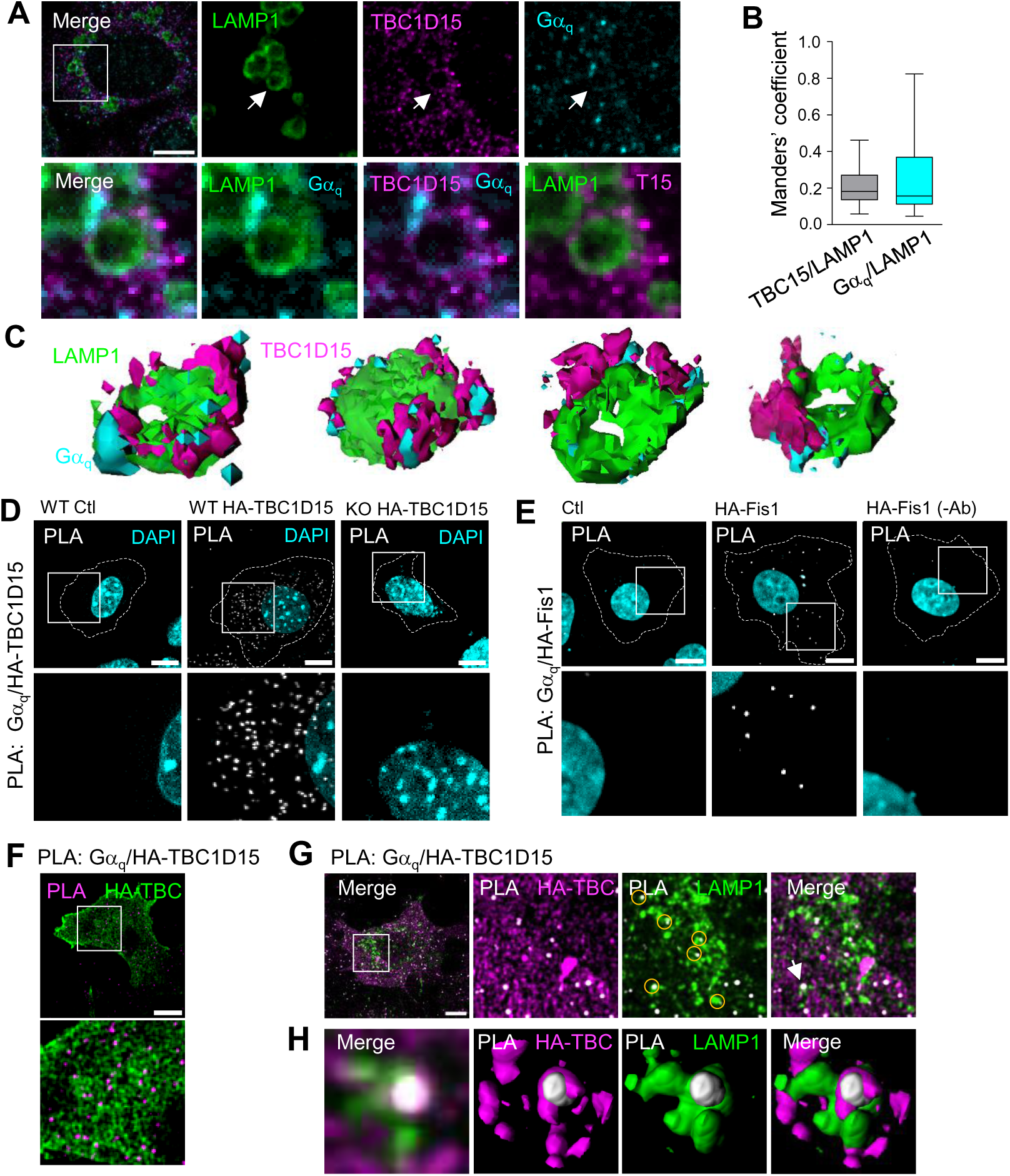
Colocalization of Ga_q_ and TBC1D15 in late endosome/lysosomes (LE/Lys). (**A**) Representative confocal microgrpahs of WT MEFs treated with U18666A (2□M, 16 h) to enlarge endolysosomes compartments and immunostained for LAMP1, TBC1D15, and Ga_q_. Inserts show enlarged regions. Arrows indicate LAMP1-positive compartments containing both TBC1D15 and Ga_q_. (**B**) Manders’ coefficients for TBC1D15/LAMP1 (TBC15/LAMP1) and Ga_q_/LAMP1 from images in (A). *n* = 22 cells. Box plots, the center line indicates the median, boxes the interquartile range, and whiskers the minimum and maximum values. (**C**) Representative 3D reconstructions of the enlarged regions shown in (A), generated using FIJI. Four examples are shown. (**D**) Representative confocal micrographs of WT and Ga_q/11_ KO (KO) MEF cells expressing HA-TBC1D15 and subjected to PLA using anti-Ga_q_ (rabbit) and anti-HA (mouse) antibodies. PLA puncta are shown in white and nuclei in cyan (DAPI). Untransfected WT cells (WT ctl) were used as a negative control. (**E**) Representative confocal micrographs of COS7 cells expressing HA-Fis1 and subjected to PLA using anti-Ga_q_ and anti-HA antibodies. Cells without primary antibodies (HA-Fis1 (-Ab)) and pcDNA3 transfected cells (Ctl) served as controls. (**F**) Representative confocal micrographs of COS7 cells expressing HA-TBC1D15 and subjected to PLA using anti-Ga_q_ and anti-HA antibodies. PLA puncta are shown in magenta and HA staining in green. (**G**) Representative confocal micrographs of COS7 cells expressing HA-TBC1D15 and LAMP1-GFP (LAMP1) and subjected to PLA using anti-Ga_q_ and anti-HA antibodies. PLA puncta are shown in white and HA staining in magenta. Enlarged regions are shown. Circles indicate PLA puncta associated with TBC1D15- and LAMP1-positive structures. (**H**) Enlarged region of the arrow indicated (G) showing PLA puncta associated with TBC1D15- and LAMP1-positive structures and 3D reconstructions within the same field of view. Dashed lines in (D) and (E) outline the cell perimeter. All scale bars are 10 µm. Images were acquired using Leica Stellaris 8 Falcon confocal microscope (A) and Andor Dragonfly microscope (D, E, F, G and H). All images are representative of at least three independent experiments.

To analyze if the Ga_q_-TBC1D15 complexes would be positioned on Lys-LE structures, cells were subjected to PLA analysis (Ga_q_-TBC1D15) in LAMP1-GFP COS7 expressing cells (Fig. 2G). Importantly, PLA puncta corresponding to Ga_q_-TBC1D15 interactions (in white) were frequently detected in close proximity to LAMP1-positive compartments (green structures) (Fig. 2G see circles in enlarged images). Furthermore, 3D reconstruction analysis facilitated the visualization of the contacts among the two proteins in Lys-LE structures, providing additional support for their spatial association (Fig 2H, S2G for different image orientations and Movie 1).

### Ga_q_ Regulation of Lysosome Size and Distribution

The localization of Ga_q_ together with TBC1D15 at endolysosomal compartments suggested a potential role for Ga_q_ in the regulation of late endosome-lysosome (LE-Lys) function and/or distribution. To investigate the impact of Ga_q_ on LE-Lys morphology, we first examined LAMP1-positive structures in wild-type (WT) and Ga_q/11_-deficient mouse embryonic fibroblasts (MEFs) (Fig. 3A-C). Ga_q/11_ knockout (KO) cells exhibited a significant increase in both the size and fluorescence intensity of LE-Lys compartments compared with WT cells. In contrast, the density of LAMP1-positive structures remained unchanged. Notably, Ga_q/11_ KO cells displayed pronounced clustering of LE-Lys compartments, with multiple lysosomes tethered together, a phenotype that was considerably less evident in WT MEFs (Fig. 3D, see detailed 3D image).

**Figure 3.**
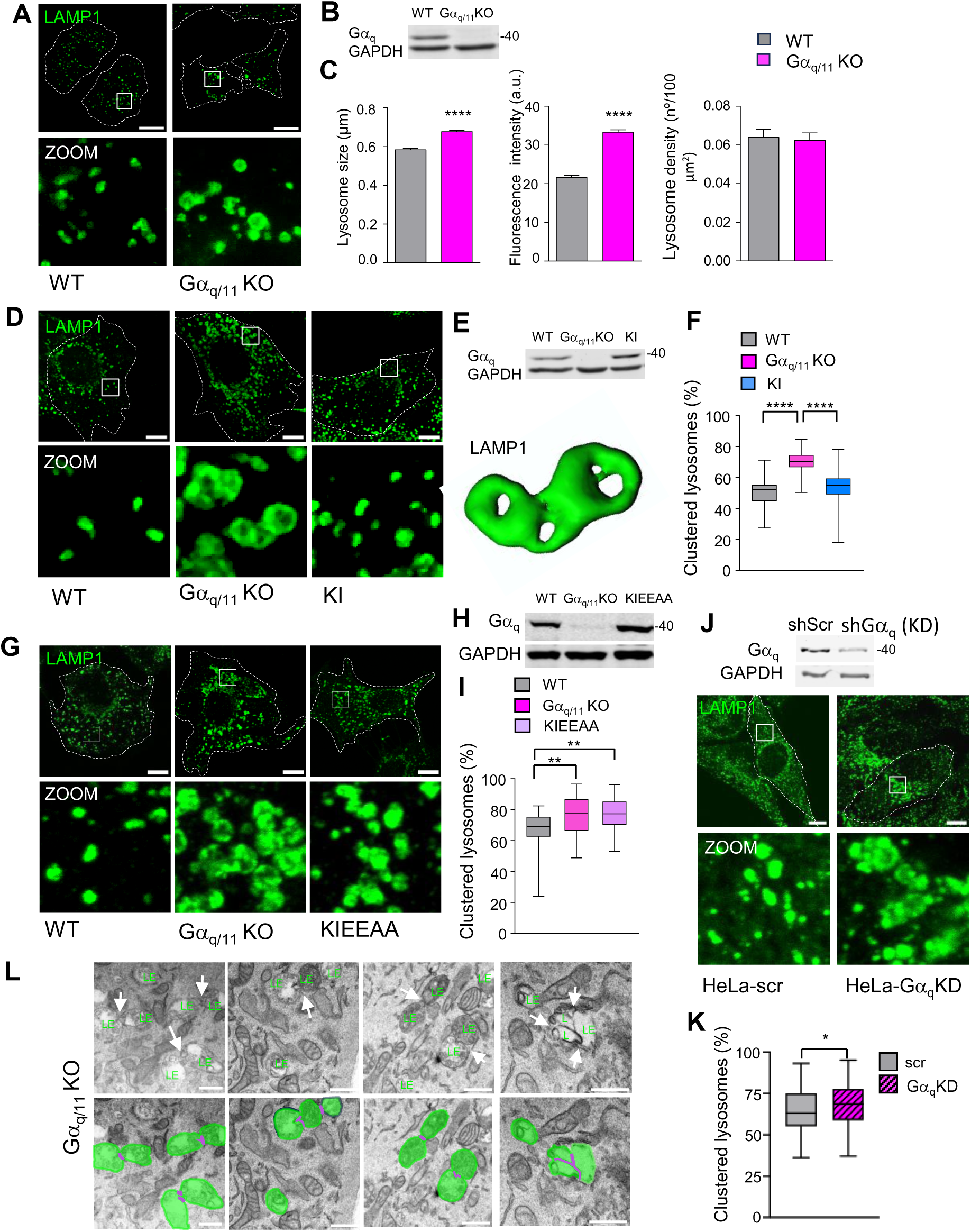
Ga_q_ regulates lysosome size and clustering. (**A**) Representative fluorescence micrographs of WT and Ga_q/11_ KO MEFs immunostained for LAMP1. Enlarged regions are shown. (**B**) Immunoblot analysis of Ga_q_ expression in WT and Ga_q/11_ KO MEFs shown in (A). Membranes were probed for Ga_q_ and GAPDH. (**C**) Quantification of lysosomal size, density (number of lysosomes per 100 µm²), and fluorescence intensity from images shown in (A) using FIJI software. WT, *n* = 31; KO, *n* = 30 cells, from 3 independent experiments. Statistics: unpaired *t*-test with Welch’s correction. (**D**) Confocal micrographs of WT, Ga_q/11_ KO, and Ga_q_ knock-in (KI) MEFs immunostained for LAMP1. 3D reconstructions of enlarged regions shown from Ga_q/11_ KO. (**E)** Immunoblot analysis of Ga_q_ expression in WT, Ga_q/11_ KO, and KI MEFs shown in (D). Membranes probed for Ga_q_ and GAPDH. (**F**) Quantification of lysosomal aggregation from (D), expressed as the percentage of lysosomal area present in clustered structures. WT, *n* = 31; Ga_q/11_ KO, *n* = 20; KI, *n* = 32 cells, from 3 independent experiments. Statistics: two-way ANOVA followed by Tukey’s multiple-comparisons test. (**G**) Confocal micrographs of WT, Ga_q/11_ KO, and Ga_q_ EEAA knock-in (KIEEAA) MEFs immunostained for LAMP1. (**H**) Immunoblot analysis of Ga_q_ expression in WT, Ga_q/11_ KO, and KIEEAA MEFs shown in (G). Membranes were probed for Ga_q_ and GAPDH. (**I**) Quantification of lysosomal aggregation from (G), expressed as the percentage of lysosomal area present in clustered structures. WT, *n* = 33; Ga_q/11_ KO, *n* = 32; KIEEAA, *n* = 46 cells, from 3 independent experiments. Statistics: Kruskal Wallis with Dunn’s multiple-comparisons test. (**J**) Immunoblot analysis of Ga_q_ expression in control (scramble) and Ga_q_-silenced (shGa_q_) HeLa cells. Membranes were probed for Ga_q_ and GAPDH. Confocal micrographs immunostained for LAMP1. (**K**) Quantification of lysosomal clustering in control and shGa_q_ (Ga_q/11_ KD) HeLa cells from images as in (J). Scr, *n* = 104; Ga_q/11_ KD, *n* = 96 cells from 3 independent experiments. Statistical significance: unpaired *t*-test with Welch’s correction. (**L**) Representative transmission electron microscopy (TEM) images of lysosomes in Ga_q/11_ KO MEFs. In all images dashed lines delineate cell boundaries. Scale bars: all images 10 µm; (A) 20 µm (M) 0.5 µm. Images were acquired using Leica AF6500 wild-field system (A), Leica Stellaris 8 Falcon confocal microscope (D, G and J) and JEOL JEM-1010 transmission electron microscope (L). Data for bar plots are mean ± SEM and for box plots, the center line indicates the median, boxes the interquartile range, and whiskers the minimum and maximum values. \**P* < 0.05, \*\**P* < 0.01, \*\*\*\**P* < 0.0001.

Importantly, these alterations were partially reversed in Ga_q/11_ KO cells stably re-expressing Ga_q_ (knock-in, KI), indicating that the observed phenotype was specifically attributable to the absence of Ga_q/11_ signaling (Fig. 3D-F). In contrast, expression of the PB1-like mutant in the Ga_q_ native form (Ga_q_EEAA) in Ga_q/11_ KO cells (KIEEAA) failed to restore the LE-Lys morphology observed in WT cells (Fig. 3G-I). This finding suggests that the regulation of lysosomal organization depends on the ability of Ga_q_ to form a functional complex with TBC1D15 and Fis1, an interaction that is disrupted by the Ga_q_EEAA mutant.

To determine whether this phenotype was cell type-specific, Ga_q_ expression was depleted in HeLa cells (Fig. 3J). Knockdown of Ga_q_ recapitulated the alterations observed in Ga_q/11_ KO MEFs (Fig. 3J and K), demonstrating that the effect is not restricted to a particular cellular background. Consistent with these observations, flat-embedded transmission electron microscopy (TEM) analysis revealed enlarged clusters of lysosomes (fake-colored green) in Ga_q/11_ KO cells (see different examples in Fig. 3L), not so frequently observed in WT cells (Fig. S3A).

### Ga_q_ Regulation of Mitochondria-Lysosome Contacts

We next investigated whether Ga_q_ influences the physical interactions between lysosomes and mitochondria. Mitochondria were visualized using TOMM20 immunostaining and analyzed together with LAMP1-positive LE-Lys compartments (Fig. 4A). Loss of Ga_q/11_ resulted in a significant increase in mitochondria-lysosome tethering events compared with WT cells (Fig. 4B, C; S4A and Movie 2 and 3). In several Ga_q/11_ KO cells, mitochondria were extensively surrounded by LAMP1-positive structures (Fig. 4A; white arrowheads), a configuration previously associated with mitochondrial fission sites. In addition, large lysosomal clusters were frequently found in close association with mitochondria in Ga_q/11_ KO cells compared to WT cells (Fig. 4B and C). A similar pattern of mitochondrial and lysosomal aggregation was observed in HeLa cells subjected to Ga_q_ knockdown, indicating that this phenotype is conserved across cell types (Fig. S4B). Similarly, close distances between lysosome and mitochondria were observed in flat-embedded TEM image analysis (Fig. 4D). Only contacts smaller than 40 nm were considered for analysis being the average contact distance between organelles in WT cells 17.7 ± 1.4 nm, consistent with published results (Xie *et al*, 2024; Yu *et al*, 2020). The quantitative analysis revealed a significant reduction in the distance between lysosomes and mitochondria in Ga_q/11_ KO cells (14.1 ± 1.1 nm) compared with WT controls (Fig. 4D and E). In some TEM images, lysosomes appear to be entirely surrounded by mitochondrial structures, supporting a close spatial association between these organelles (Fig. 4F, see scheme). TEM images of KI cells suggested the restoration of contacts (Fig. 4SC). In fact, re-expression of Ga_q_ (KI) restored the number of mitochondria-lysosome contacts to WT levels (Fig. 4G and H), confirming that the observed phenotype was specifically caused by the loss of Ga_q/11_. In contrast, stable expression of the PB1-like Ga_q_EEAA mutant failed to normalize the number of contacts (Fig. 4I and J), indicating that regulation of mitochondria-lysosome contacts requires the non-canonical functions of Ga_q_. To directly assess the involvement of mitochondrial protein Fis1, Fis1 expression was reduced in Ga_q/11_ KO cells (Ga_q/11_ KO-Fis1KD) (Fig. 4K, L and S4D). Remarkably, Fis1 knockdown restored mitochondria-lysosome tethering to levels comparable to those observed in control cells.

**Figure 4.**
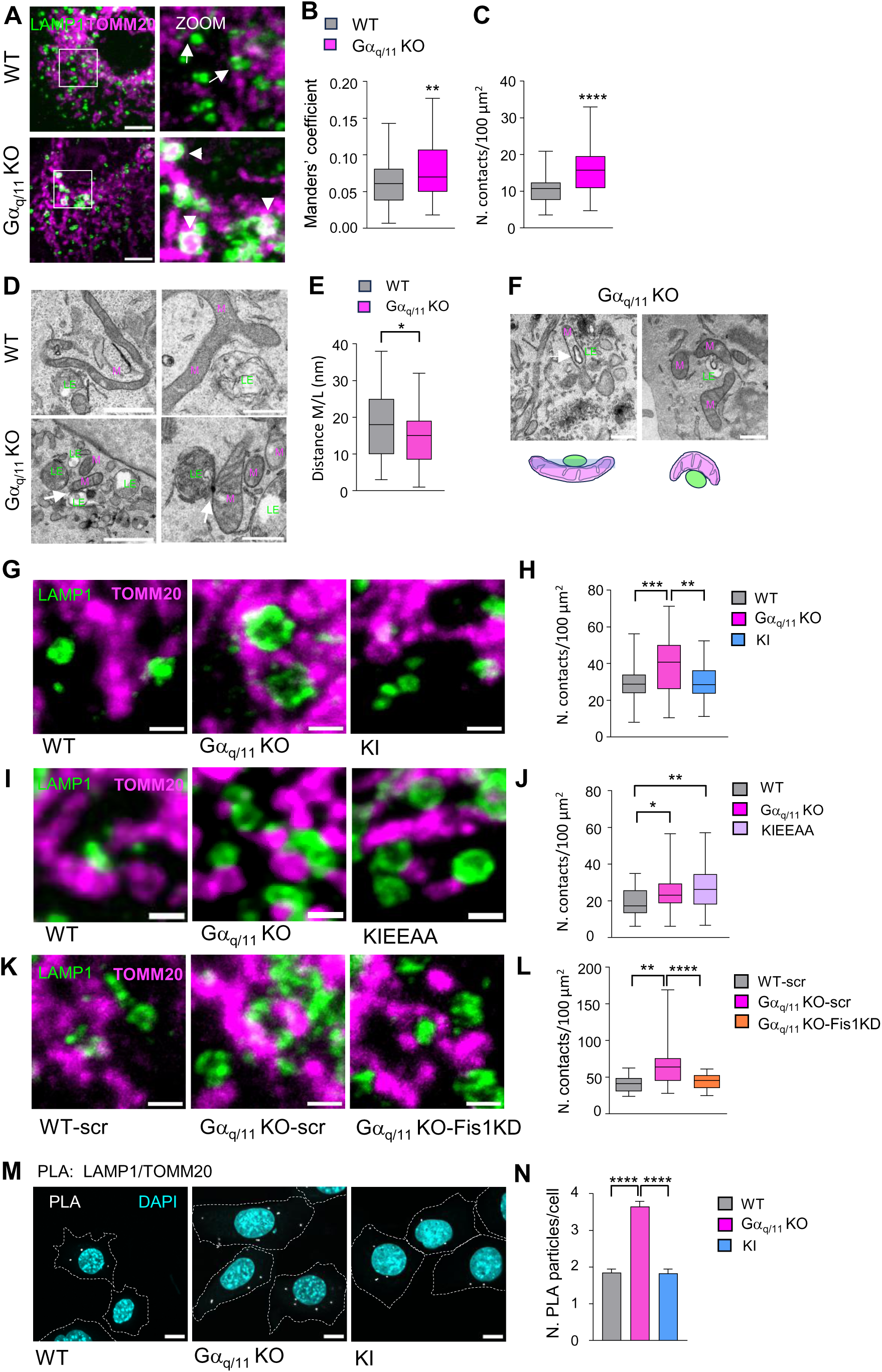
Increased contacts between LE-Lys and mitochondria in absence of Ga_q_. (**A**) Representative confocal micrographs of WT and Ga_q/11_ KO MEFs immunostained for LAMP1 and TOMM20. Arrows indicate close apposition between LAMP1 and TOMM20, whereas arrowheads denote mitochondria surrounded by LAMP1-positive structures. Scale bar, 5 µm. (**B, C**) Manders’ colocalization coefficient and number of LAMP1-TOMM20 contacts per 100 µm² from images in (A) from *n* = 60 cells from three independent experiments. (**D**) Representative TEM images showing LE-Lys (LE) in close proximity to mitochondria (M) in WT and Ga_q/11_ KO MEFs. Scale bar: 0.5 µm. (**E**) Quantification of the distance between mitochondria and LE-Lys from (D) from WT, *n* = 47; Ga_q/11_ KO, *n* = 44 contacts from 2 independent experiments. (**F**) TEM images of Ga_q/11_ KO MEF cells also show endolysosomes/lysosomes (LE) embedded within mitochondrial (M) structures; a schematic representation of the contact site is included. Scale bar, 0.5 µm. (**G, I** and **K**) Enlarged confocal micrographs of WT, Ga_q/11_ KO, and KI MEFs (G); WT, Ga_q/11_ KO, and KIEEAA MEFs (I); and WT-scr, Ga_q/11_ KO-scr, and Ga_q/11_ KO-Fis1KD MEFs (K) immunostained for LAMP1 and TOMM20. Scale bar, 1 µm. (**H**, **J** and **L**) Quantification of contact numbers per 100 µm² is shown in G, I and K (H, J and L, respectively). For **H**: WT, *n* = 41; Ga_q/11_ KO, *n* = 39; KI, *n* = 39; **J**: WT, *n* = 46; Ga_q/11_ KO, *n* = 55; KIEEAA, *n* = 47; and **L**: WT, *n* = 31; Ga_q/11_ KO-scr, *n* = 29; Ga_q/11_ KO-Fis1KD, *n* = 24 cells from 2-3 independent experiments done in duplicate. Scale bar, 1 µm. (**M**) Confocal micrographs of WT and Ga_q/11_ KO cells subjected to PLA using anti-LAMP1 and anti-TOMM20 antibodies (white) showing DAPI (cyan) staining. Scale bar, 10 µm. (**N**) Quantificatioon of PLA puncta per cell from (M) from more than *n* = 100 cells from 3 independent conditions. Images were acquired using Leica Stellaris 8 Falcon confocal microscope (A, G, I and K), JEOL JEM-1010 transmission electron microscope (D and F) and Andor Dragonfly microscope (M). Data for bar plots are mean ± SEM and for box plots, the center line indicates the median, boxes the interquartile range, and whiskers the minimum and maximum values. Statistics: Unpaired t-test (B) with Welch’s correction (C and E), One-way ANOVA with Turkey’s multiple comparison test (H) and Kruskal Wallis with Dunn’s multiple-comparisons test (J, L and N). \**P* < 0.05, \*\**P* < 0.01, \*\*\**P* < 0.001, \*\*\*\**P* < 0.0001.

To independently validate these findings, mitochondria-lysosome interactions were quantified using PLA with antibodies against LAMP1 and TOMM20. Ga_q/11_ KO cells exhibited a significant increase in PLA signal relative to WT cells, indicative of enhanced proximity between lysosomes and mitochondria (Fig. 4M and N). Re-expression of Ga_q_ reduced the number of PLA puncta to WT levels, whereas no signal was detected in the corresponding negative controls (Fig. S4E).

### Ga_q_ Regulates Mitochondria-Lysosome contact dynamics

The absence of Ga_q/11_ results in lysosomal enlargement, clustering, and increased mitochondria-lysosome contacts. However, the elevated number of contacts observed in Ga_q/11_ KO cells could potentially be explained by the altered distribution and clustering of lysosomes. To determine whether Ga_q_ directly regulates organelle tethering dynamics, we performed live-cell time-lapse microscopy to monitor interactions between LE-Lys compartments and mitochondria (Fig. 5; Movies 4, 5, 6 and 7).

**Figure 5.**
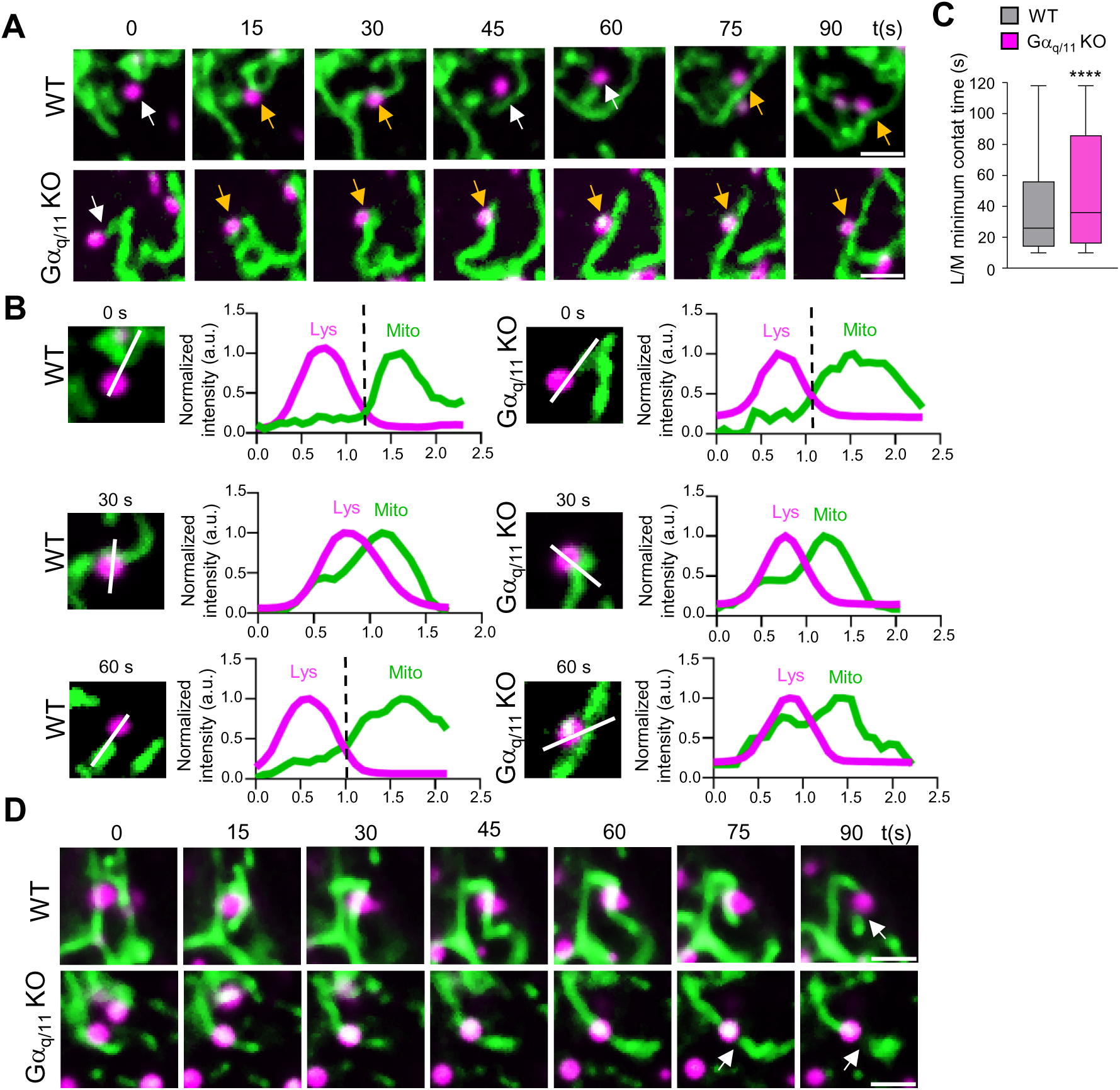
(**A**) Representative confocal micrographs of WT and Ga_q/11_ KO cells stained with LysoTracker (magenta) and MitoTracker (green). White arrows indicate separated lysosomes and mitochondria, whereas yellow arrows denote lysosome-mitochondria contacts. (**B**) Magnified images at 0, 30 and 60 s for each cell type. Corresponding normalized LysoTracker (magenta) and MitoTracker (green) fluorescence intensity profiles along the indicated white line are shown. (**C**) Minimum duration of lysosome-mitochondria (L/M) contacts (s) from the experiments shown in A with more than 100 contacts analyzed, from 3 idependent experiments. In box plots, the center line indicates the median, boxes the interquartile range, and whiskers the minimum and maximum values. Statistics: Mann-Whitney test. **** *P* < 0.0001. (**D**) Representative confocal micrographs of WT and Ga_q/11_ KO cells stained with LysoTracker (magenta) and MitoTracker (green). White arrows indicate mitochondria fission. All scale bars, 2 µm. All images were acquired using Andor Dragonfly microscope.

Mitochondria and lysosomes were visualized using MitoTracker Green and LysoTracker Deep Red, respectively. In WT cells, most mitochondria-lysosome contacts were highly transient, typically lasting around 20 s. These interactions displayed a characteristic “kiss-and-run” behavior, in which lysosomes rapidly associated with and detached from mitochondria without following a defined trajectory, appearing to move in a discontinuous or hopping manner (Fig. 5A; see normalized line intensity profiles 5B; Movie 4). This behavior is consistent with the dynamic and reversible nature of mitochondria-lysosome contacts previously described (Wong *et al*, 2018). In contrast, Ga_q/11_ KO cells exhibited a markedly different interaction profile. A substantial fraction of lysosomes compartments remained associated with mitochondria for prolonged periods, frequently exceeding 20 s in duration (Fig. 5C, Movie 5). In many instances, lysosomes traveled along mitochondrial tubules while maintaining continuous contact, indicating a failure in the normal detachment process. These trajectories frequently resembled a “walking” movement of lysosomes along the mitochondrial network (Fig. S5A, arrows), suggesting enhanced tethering between the two organelles.

Despite these differences in contact dynamics, mitochondria-lysosome interactions in both WT and Ga_q/11_ KO cells were occasionally followed by mitochondrial fragmentation events (Fig. 5D, arrows; Movies 6 and 7), consistent with the proposed role of lysosome-associated contact sites in mitochondrial fission. In addition, a subset of lysosomes remained static in both genotypes, indicating the coexistence of stable and dynamic populations of organelle contacts. Importantly, the prolonged interactions observed in Ga_q/11_ KO cells demonstrate that the increase in mitochondria-lysosome contacts cannot be explained solely by altered lysosomal positioning or clustering. Rather, the absence of Ga_q/11_ directly enhances organelle tethering by impairing the mechanisms that normally promote contact dissociation.

Collectively, these findings support a model in which Ga_q_ functions as a critical regulator of mitochondria-lysosome contact dynamics. Through a mechanism dependent on Fis1, Ga_q/11_ regulate contact dissociation between LE-Lys compartments and mitochondria.

### Fis1 and TBC1D15 interactions with Ga_q_ Regulates Mitochondrial Fission

One of the key functional consequences of mitochondria-lysosome contacts is the promotion of mitochondrial fission, which frequently occurs at sites where lysosomes transiently interact with the mitochondrial network (Wong *et al*, 2018; Kleele *et al*, 2021). Analysis of TEM images from Ga_q/11_ knockout (KO) revealed the presence of lysosomes in close proximity to mitochondrial fission sites located at the extremities of mitochondria and often associated with endoplasmic reticulum (ER) (Boutry & Kim, 2021) (Fig. 6A). Mitochondria-lysosome contacts in sites of mitochondrial fragmentation were also observed in WT cells.

**Figure 6.**
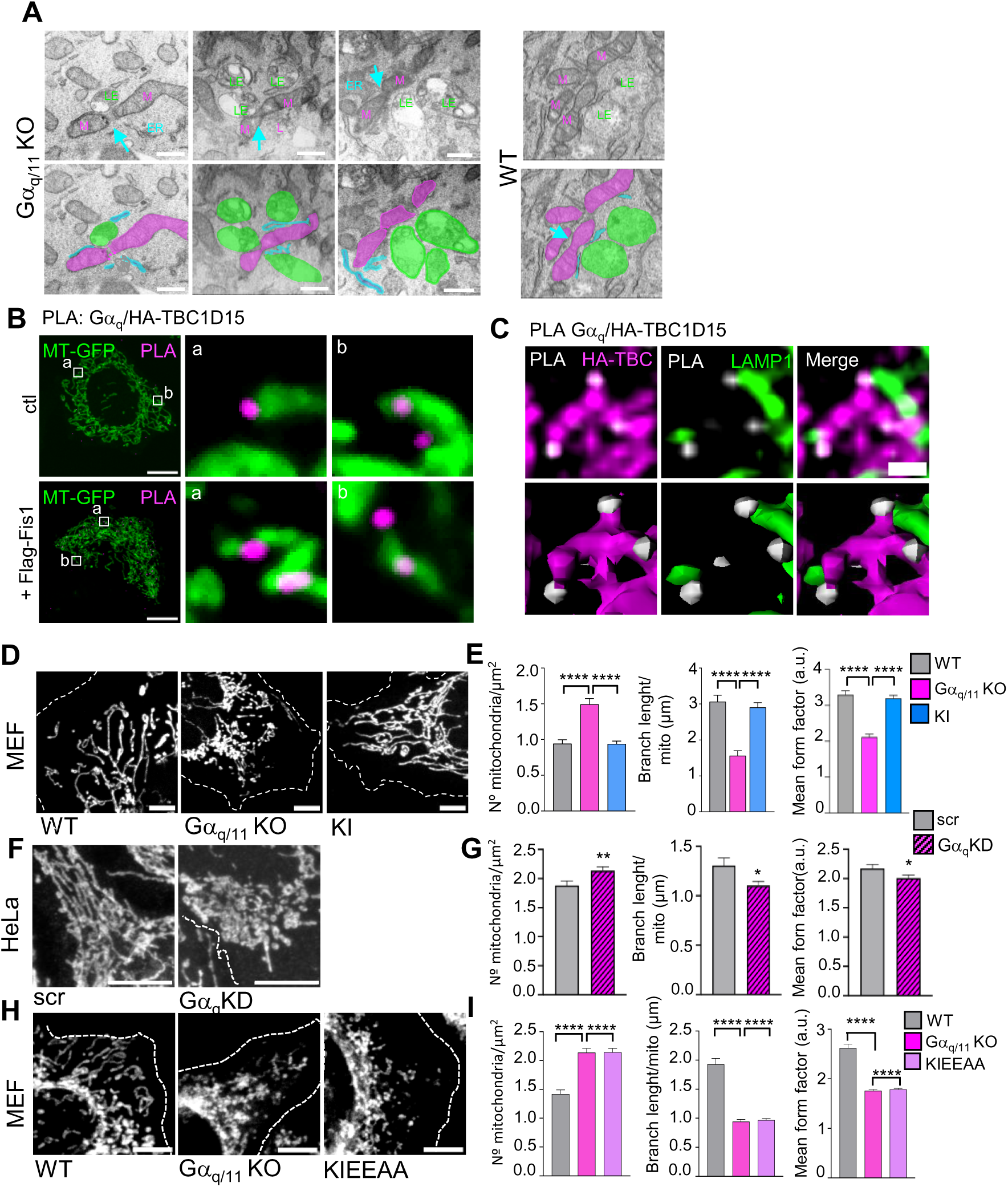
Ga_q_ regulates mitochondrial fission. (**A**) Three representative TEM images from Ga_q/11_ KO and one WT showing late endosome-lysosomes (LE-Lys; LE) in close proximity to mitochondrial (M) fission sites in MEFs. Pseudocolor overlays show LE-Lys (green), mitochondria (magenta), and ER (cyan). Scale bar 0.5 µm. (**B**) Representative confocal micrographs of WT MEFs cells expressing MT-GFP (mitochondria), HA-TBC1D15 and/or Flag-Fis1 and subjected to PLA using anti-Ga_q_ and anti-HA antibodies (magenta). Two enlarged images are shown. Scale bar, 10 µm. (**C**) Representative confocal micrographs of COS7 cells expressing Flag-Fis1 and HA-TBC1D15. PLA puncta (white), indicating Ga_q_-HA-TBC1D15 interactions, are detected in distal TBC1D15-positive structures (magenta) adjacent to LAMP1-positive compartments (green). Enlarged regions are shown. Scale bar, 1 µm. (**D**, **F** and **H**) Representative enlarged confocal micrographs of WT, Ga_q/11_ KO, and KI MEFs (**D**); HeLa-scr and HeLa-Ga_q_KD cells (**F**); and WT, Ga_q/11_ KO, and KIEEAA MEFs (**H**) showing mitochondria stained with MitoTracker Red CMXRos (white). Scale bar, 5 µm. (**E**, **G**, **I**) Quantification of mitochondrial morphology, including mean form factor, branch length/mitochondria, and mitochondrial number/µm^2^, from images shown in (D), (F) and (H), respectively. Data were obtained from *n = 3* independent experiments and are presented as mean ± SEM. For (**E**): WT, *n* = 65, Ga_q/11_ KO, *n* = 56, and KI, *n* = 59; (**G**) Scr, *n* = 46; Ga_q/11_ KD, *n* = 64 cells; (**I**) WT, *n*=57; Ga_q/11_ KO, *n*=52; KIEEAA, *n*= 64 cells from 3 independent experiments. Statistics: Kruskal Wallis with Dunn’s multiple-comparisons test (**E** and **I**) and Mann-Whitney test (G). Images were acquired using a JEOL JEM-1010 transmission electron microscope (A), a Leica Stellaris 8 Falcon microscope (B, D, F and H) and an Andor Dragonfly microscope (C). \**P* < 0.05, \*\**P* < 0.01, \*\*\*\**P* < 0.0001.

To determine whether the Ga_q_-TBC1D15 complex localizes to these fission-associated contact sites, we analyzed the distribution of Ga_q_-TBC1D15 interactions by PLA in MEF cells that expressed Mito-GFP (MT-GFP), which exclusively localizes at the mitochondria (Fig. 6B, a,b and S6A). Most of PLA puncta, reflecting Ga_q_-TBC1D15 interactions, where located close to mitochondria and specifically at the edges of mitochondrial structures. To facilitate the localization of TBC1D15 at the mitochondria, we utilized cells that over-expressed Fis1 (Fig. S6B and C) (Onoue *et al*, 2013; Yamano *et al*, 2014). In these conditions TBC1D15 localized predominantly at mitochondria (Onoue *et al*, 2013; Yamano *et al*, 2014). As before, PLA puncta corresponding to Ga_q_-TBC1D15 interactions were frequently detected at the periphery of mitochondrial structures. Some of the signals localized in locations that resemble mitochondrial fission sites (Fig. 6B and S6C). Further, we analyze the position of Ga_q_-TBC1D15 interactions in COS7 cells expressing LAMP1-GFP in the presence of Fis1 (Fig. S6D). Enlarge confocal images show PLA signals positioned adjacent to TBC1D15-enriched-mitochondria and close to LAMP1 structures (Fig. 6C, S6D and Movie 8). As before these PLA puncta where predominantly located at the edges of the TBC1D15-enriched-mitochondria. Together these results suggest that the Ga_q_-TBC1D15 complex localizes in close proximity to locations where lysosome-associated mitochondrial fission is likely to occur.

To evaluate the functional consequences of Ga_q_ depletion on mitochondrial morphology, WT, Ga_q/11_ knockout (KO), and KI MEFs were stained with MitoTracker Red ROS to visualize the mitochondrial network. Loss of Ga_q/11_ resulted in a marked increase in mitochondrial fragmentation, whereas re-expression of Ga_q_ restored mitochondrial morphology to a pattern resembling that of WT cells (Fig. 6D and E). Mitochondria in Ga_q/11_ KO cells appeared shorter and more punctate, indicative of enhanced fission activity. In contrast, KI cells displayed a more interconnected and elongated mitochondrial network, supporting a role for Ga_q_ in maintaining mitochondrial integrity. These results were corroborated in HeLa Ga_q_ depleted cells (Fig. 6F and G). Interestingly, stable expression of the PB1-like mutant Ga_q_EEAA in MEFs resulted in an even more pronounced fragmented phenotype, accompanied by the accumulation of mitochondria in the perinuclear region (Fig. 6H and I). Quantitative analysis of mitochondrial morphology in Fis1-KD cells promoted mitochondrial elongation and reducing fragmentation as shown before (Stojanovski *et al*, 2004) and contrary to the Ga_q/11_ KO phenotype. Remarkably, simultaneous depletion of Ga_q/11_ and Fis1 restored mitochondrial morphology to levels comparable to those observed in WT control cells (Fig. S6E).

These findings support a model in which Ga_q_ acts as a regulator of mitochondria-lysosome untethering. In the absence of Ga_q_, increased and prolonged contacts between lysosomes and mitochondria promote Fis1-dependent mitochondrial fission, resulting in a fragmented mitochondrial network.

### Ga_q_ Regulates Mitophagy Through Fis1

Mitochondrial fission is a prerequisite for efficient mitophagy, and lysosome-associated fission events have been proposed to facilitate the delivery of mitochondrial fragments to the autophagic pathway. In addition, TBC1D15 has been implicated in the regulation of mitophagy through its interactions with Fis1, Rab7, and LC3 proteins. TBC1D15 contains conserved LC3-interacting region (LIR) domains, which are crucial for proper LC3 orientation and isolation membrane growth (Yamano *et al*, 2014). Previous transmission electron microscopy (TEM) analyses revealed an increased abundance of autophagosomes and autolysosomal structures in Ga_q/11_ knockout (KO) cells (Benincá *et al*, 2014). Here, we further investigated the impact of Ga_q/11_ depletion on mitophagy induced by valinomycin.

To monitor mitophagy, we employed the dual-fluorescent reporter Mito-QC. Under basal conditions, mitolysosomes were rarely detected in wild-type (WT) cells (Fig. 7A and B). In contrast, Ga_q/11_ KO cells displayed a substantial number of rounded mitolysosomal structures even in the absence of valinomycin treatment. Upon exposure to valinomycin (3 h), the number of mitolysosomes increased significantly in Ga_q/11_ KO cells, exceeding the levels observed in treated WT controls. Consistent with these findings, downregulation of Ga_q_ in sh-treated HeLa cells stably expressing Parkin-YFP (Fig. S7A and B) resulted in a marked accumulation of LC3-RFP-positive puncta associated with Parkin-labelled mitochondria following valinomycin treatment (Fig. S7C). Notably, loss of Ga_q_ did not impair Parkin recruitment to mitochondria, indicating that Ga_q_ does not affect the early stages of mitophagy involving Parkin translocation and mitochondrial outer membrane ubiquitination. Together, these observations suggest that Ga_q/11_ deficiency enhances mitophagic activity.

**Figure 7.**
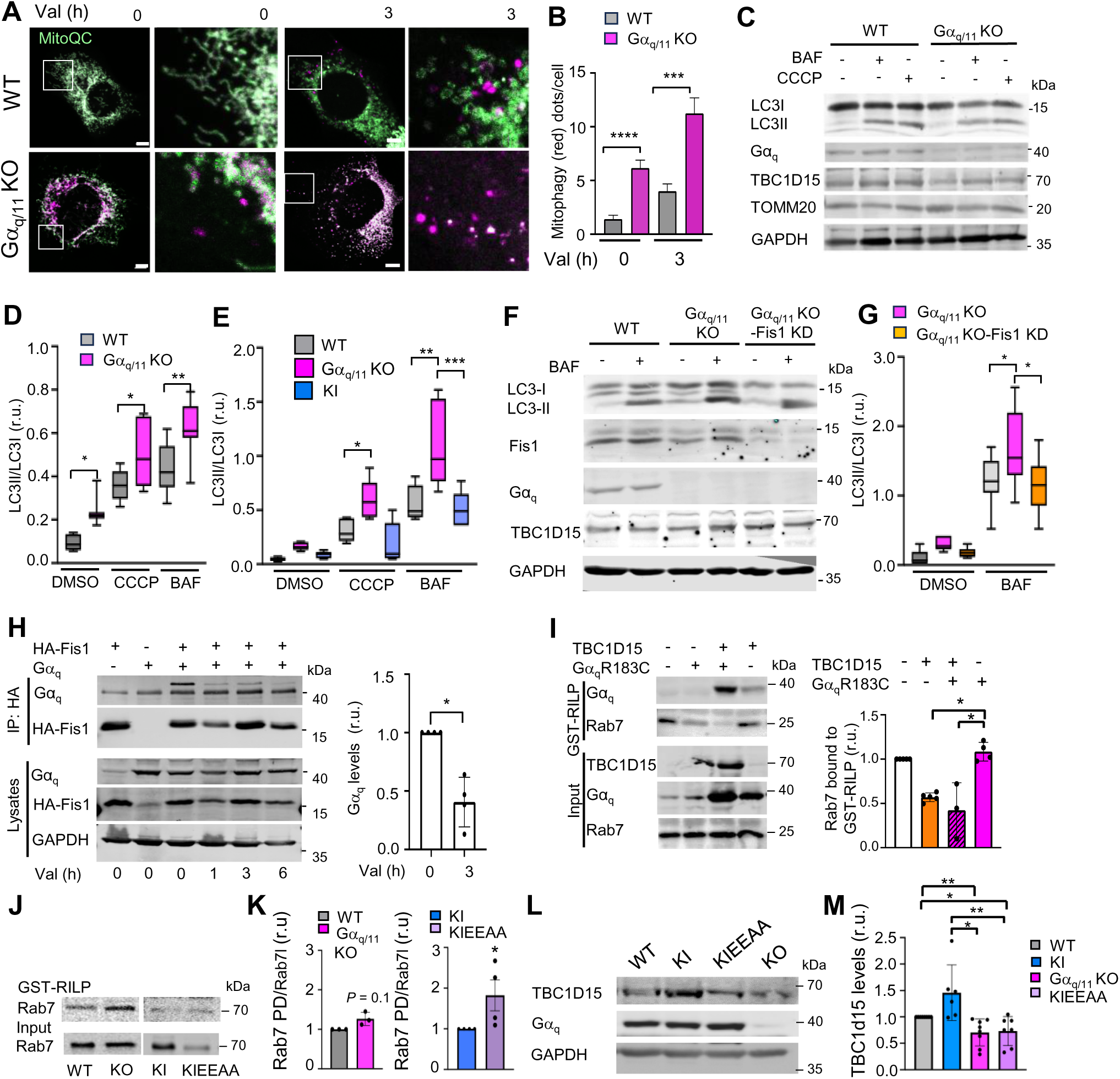
Increased mitophagy in absence of Ga_q/11_ dependent of Fis1 interaction. (**A**) Confocal micographs of WT and Ga_q/11_ KO cells expressing mito-QC and myc-parkin treated with valinomycin (10 mM) for 0 and 3 h. Insets show image magnification. Images were acquired using a LSM700 confocal microscope (Zeiss). Scale bar 5 µm. (**B**) Quantification of mitophagy from images as in (A). Experiments were done in *n* = 4 independent experiments. *n* = 56 (WT-0 h Val), 62 (Ga_q/11_ KO-0 h Val), 65 (WT-3 h Val), 77 (Ga_q/11_ KO-3 h Val) cells. (**C**) Representative immunoblots from WT and Ga_q/11_ KO MEFs treated with CCCP (10 mM) for 3 h or with bafilomycin (BAF) (100 nM) for 5 h. Samples were immunoblotted with antibodies against Ga_q_, LC3, TOMM20, GAPDH and TBC1D15. (**D**) LC3II/LC3I ratio quantification from (C) from n=8 samples each condition from *n = 4* independent experiments. (**E**) LC3II/LC3I ratio of WT, Ga_q/11_ KO and KI MEF cells treated with CCCP (10 mM) for 3 h or with bafilomycin (BAF) (100 nM) for 5 h from gels as in (Fig. 7SE). *n = 5* independent experiments. (**F**) Representative immunoblots from WT, Ga_q/11_ KO and Ga_q/11_ KO/Fis1KD MEFs treated with bafilomycin (BAF) (100 nM) for 5 h. Samples were immunoblotted with antibodies against Ga_q_, LC3, GAPDH, Fis1 and TBC1D15. (**G**) LC3II/LC3I ratio from gels as in (F) from *n = 2* independent experiments done in duplicated. (**H**) HeLa-parkin-YFP cells expressing Ga_q_ and HA-Fis1 were treated with Val (10 mM) for the times indicated before immunoprecipitation with antibodies against anti-HA. Samples were immunoblotted with antibodies against HA, Ga_q_ and GAPDH. Quantification of Ga_q_ levels from *n = 4* independent experiments. Samples were normalized by GAPDH and relative to Ga_q_ immunoprecipitation. (**I**) GST-RILP pull-down (PD) assay form HEK293 cells expressing HA-TBC1D15 in the presence or absence of Ga_q_R183C, immunostained against Rab7, Ga_q_ and TBC1D15. Quantification of bands from a minimum of three replicates. Data are means ± SEM. (**J**) GST-RILP PD assay from WT, Ga_q/11_ KO, KI, KIEEAA MEF cell extracts, immunostained to Rab7. Input sample is shown. (**K**) Quantification of bands as in J from 3 (WT, Ga_q/11_ KO) and 4 (KI, KIEEAA) replicates. (**L**) Representative immunolots form WT, Ga_q/11_ KO, KI, KIEEAA MEF cells immunoblotted for TBC1D15, Ga_q_ and GAPDH. (**M**) Quantification of bands as in (M) from *n = 7* experiments. Data for bar plots are mean ± SEM and for box plots, the center line indicates the median, boxes the interquartile range, and whiskers the minimum and maximum values. Statistics: Kruskal-Wallis with Dunn’s multiple comparison test (B, D, K, N), One way Anova with Tukey (E, G) and Mann-Whitney test (H, J). \**P* < 0.05, \*\**P* < 0.01, \*\*\**P* < 0.001, \*\*\*\**P* < 0.0001.

To further assess autophagic flux, we evaluated LC3 lipidation. Under basal conditions, Ga_q/11_ KO cells exhibited a higher LC3-II/LC3-I ratio than WT cells (Fig. 7C and D), indicative of increased autophagic activity. Chronic treatment with the mitochondrial uncoupler carbonyl cyanide m-chlorophenyl hydrazone (CCCP) further increased LC3 lipidation in Ga_q/11_ KO cells relative to vehicle-treated controls. Importantly, inhibition of lysosomal degradation with bafilomycin A1 resulted in an additional accumulation of LC3-II, supporting the notion that the absence of Ga_q/11_ enhances autophagic flux rather than merely impairing autophagosome turnover. Effect that was not present in KI recovered cells further supporting the specific effect of Ga_q/11_ absence (Fig. 7E and S7D). To determine whether this phenotype requires Fis1, LC3 lipidation was analyzed in Ga_q_-deficient MEFs subjected to Fis1 knockdown (Fig. 7F and G). Fis1 depletion significantly reduced the LC3-II/LC3-I ratio in Ga_q_ KO cells, indicating that the elevated autophagic flux associated with Ga_q_ loss is dependent on Fis1. These results identify Fis1 as a key mediator of the autophagic phenotype observed in the absence of Ga_q_.

We next investigated whether mitophagy induction influences the interaction between Ga_q_ and Fis1. Co-immunoprecipitation experiments in HeLa-Parkin cells revealed that valinomycin treatment (1, 3 and 6 h) significantly reduced the amount of Ga_q_ associated with Fis1 (Fig. 7H). These findings suggest that dissociation of Ga_q_ from the Fis1-TBC1D15 complex may be required to promote mitophagy, supporting a model in which Ga_q_ regulates the availability of TBC1D15 for interaction with Fis1 and thereby restrains excessive mitophagic activity. Interestingly, mitophagy induction did not decrease the interaction between Fis1 and constitutively active Ga_q_ (Fig. S7E), suggesting that the activation state of Ga_q_ may influence the dynamics of complex disassembly during mitophagy.

### Ga_q_ regulates Rab7 activity through TBC1D15

Rab7 activity is essential for the regulation of mitochondria-lysosome contacts, in addition to its well-established role in mitophagy. We therefore evaluated Rab7 activity in Ga_q/11_ KO cells. To do so, we took advantage of the ability of active Rab7 to bind its effector protein RILP. As expected, the constitutively active Rab7 mutant efficiently bound the GST-RILP fragment, whereas the dominant-negative mutant did not (Fig. S7F). Also, TBC1D15 expression reduced Rab7 binding to GST-RILP consistent with its GAP activity (Fig. 7I). Cell extracts from Ga_q/11_ KO cells displayed increased Rab7 activity compared with WT cells, and this increase was higher in KI cells expressing the PB1-like domain mutant, Ga_q_EEAA (KIEEAA) (Fig. 7J and K). But, expression of Ga_q_ alone did not affect the Rab7 activity or its protein levels (Fig. 7I).

Interestingly, TBC1D15 protein levels were reduced in both Ga_q/11_ KO and KIEEAA cells which could explain the increase Rab7 activity (Fig. 7L and M). These findings suggest that Ga_q_ may contribute to the stabilization of TBC1D15, thereby maintaining appropriate cellular levels of this protein. Therefore, the presence of Ga_q_ appears to be required for the proper regulation of the Rab7-associated molecular complex. In this context, we found that the active, GTP-bound form of Rab7 co-immunoprecipitated with Ga_q_ (Fig. S7G), indicating that active Rab7 forms a complex with both TBC1D15 and Ga_q_ more likely in the organelle contacts.

Taken together, these findings support a model in which Ga_q_ forms a complex with Fis1 and TBC1D15 that regulates the stability of the complex and the activity of Rab7 which in turn coordinates mitochondria-lysosome contacts, mitochondrial fission, and mitophagy. Disruption of this complex in the absence of Ga_q_ enhances Rab7 activity and in turn organelle contacts, and stimulates mitophagic flux, whereas regulated dissociation of Ga_q_ from Fis1 appears necessary to facilitate mitochondrial turnover under mitophagy-inducing conditions (Fig. 8).

**Figure 8.**
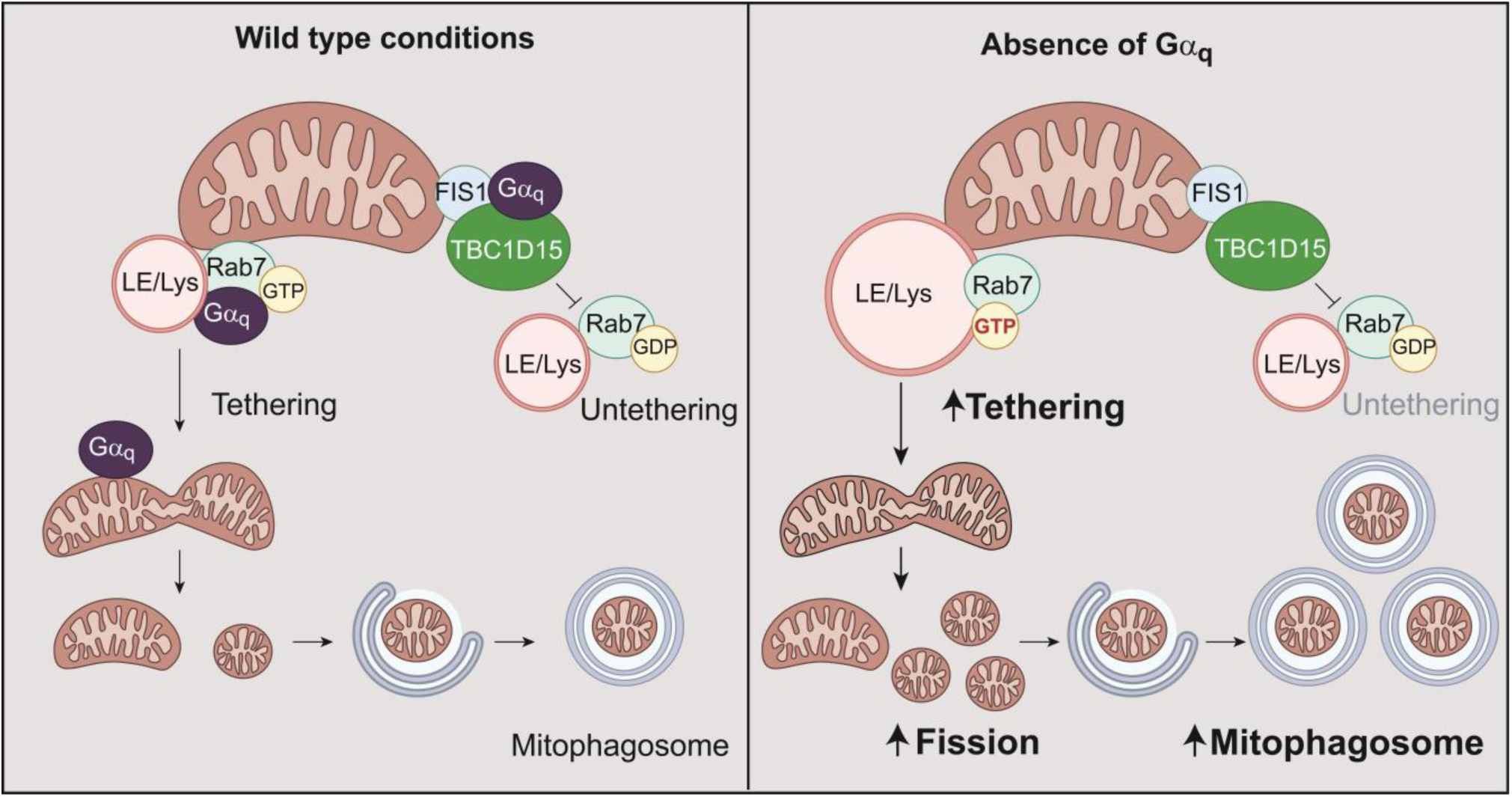
Model of Ga_q_ acting as a ‘tether-breaker’, promoting the release of contacts. The tethering of LE/Lys and mitochondria (M) is regulated by the activity of Rab7-GTP. The complex formed by Fis1 and Ga_q_ facilitates the untethering process by forming a complex with TBC1D15 that, in turn, inhibits the activity of Rab7. The tethering leads to an asymmetric fission of mitochondria. The smaller daughter is engulfed inside the mitophagosome. In wild type conditions there is a balance between tethering and untethering (left side). In absence of Ga_q_ there is an increased tethering, leading to bigger LE/Lys, more fission of the mitochondria and a higher number of mitophagosomes (right side). Model created with Adobe Illustrator.

## Discussion

Lysosomes and mitochondria are highly dynamic organelles that are essential for cellular homeostasis, and dysfunction of either organelle has been implicated in numerous diseases (Kiraly *et al*, 2025; Liu *et al*, 2026). Recent advances have demonstrated that contact sites between these two organelles play important roles in regulating their cellular distribution and function (Wong *et al*, 2022). Understanding how these contacts are regulated is therefore critical for understanding their contribution to cellular physiology. Here, we provide evidence for a role of Ga_q_ in maintaining the number and duration of lysosome-mitochondria contacts through regulation of the Fis1-TBC1D15-Rab7 pathway. We propose that Ga_q_ acts as a "tether breaker" that controls the balance between contact formation and dissociation. This work establishes a novel link between intracellular G protein signaling and inter-organelle communication.

The present findings demonstrate that Ga_q_ is physically linked to the molecular machinery governing mitochondrial fission and mitophagy through a non-canonical signaling pathway. Notably, the non-canonical effector region of Ga_q_ has previously been implicated in interactions with several intracellular proteins, including PKCζ and p62, which have been associated with the regulation of autophagy (Cabezudo *et al*, 2021; Sánchez-Fernández *et al*, 2013, 2016). Together, with the present results, these observations support the existence of a novel signaling axis that cooperates with canonical GPCR pathways to regulate organelle homeostasis. More broadly, our findings highlight the non-canonical intra-organelle signaling pathway as an additional pathway that contributes to important physiological functions within the cell.

Fis1 is known to promote mitochondrial fission through its interactions with Mfn1, Mfn2, Mid51, and Drp1, besides its documented association to TBC1D15 and TBC1D17 (Shi *et al*, 2023; Wong *et al*, 2022). Here, we provide evidence for a previously unrecognized interaction between Ga_q_ and the Fis1 TPR-domain (Egner *et al*, 2022), located in proximity to the TBC1D15-binding region (Onoue *et al*, 2013), and consistent with the formation of a ternary complex. Under physiological conditions, mitophagy induction promotes the dissociation of Ga_q_ from Fis1, thereby enabling Fis1 to engage additional binding partners required for mitochondrial fission and/or autophagosome formation (Figure 8). Consistent with this model, Ga_q_ absence leads to increased mitochondrial fission and mitophagy, suggesting that Ga_q_ acts as a negative regulator that restrains these processes under basal conditions. Interestingly, the N-terminal region of Fis1, located upstream of the TPR domain, has been implicated in Drp1 binding (Egner *et al*, 2022; Ihenacho *et al*, 2023). In parallel, mitophagy induction requires the oligomerization of Fis1 and TBC1D15 (and/or TBC1D17), facilitating their interaction with the autophagosomal membrane through the LIR motifs present in TBC1D15 (Yamano *et al*, 2014). Notably, several of these LIR motifs are located in close proximity to the Ga_q_ and Fis1 binding region, raising the possibility that Ga_q_ binding may influence the assembly or remodeling of protein complexes involved in coordinating mitochondrial fission and mitophagy.

Recent work has provided evidence that the division of mitochondria through Drp1 proteins at the periphery enables mitochondria to be shared into smaller mitochondria destined for mitophagy is regulated by lysosome and Fis1 (Kleele *et al*, 2021). Midzone fission leads to proliferation of mitochondria and requires ER and actin structures. Peripheral fission is facilitated by lysosomal contacts regulating mitochondria asymmetric division destined for mitophagy (Kleele *et al*, 2021). Lysosomes are present at approximately 60% of the sites of mitochondria division, where ER is also recruited (Boutry & Kim, 2021). Compatible with the present results, we had previously found that Ga_q/11_ overexpression induces hyperfused mitochondria in a mechanism that is partially independent of Mfn1/2 (Benincá *et al*, 2014; Izquierdo-Villalba *et al*, 2024). Also we had observe that Ga_q/11_ localizes in mitochondria in scattered punctate areas (Izquierdo-Villalba *et al*, 2024). In the present study, we show Ga_q_ positioned at the periphery of mitochondria in close contact to TBC1D15 and to endolysosome structures. In absence of Ga_q/11_ contacts persist for longer periods of time compared with wild-type cells. Ga_q/11_ effects prevent excessive stabilization of mitochondria-lysosome associations which in turn decreases mitochondria fragmentation and mitophagy. Therefore, Ga_q/11_ is necessary at the contact sites between mitochondria and lysosomes to regulate the Fis1-TBC1D15-Rab7 axis most likely targeted to mitophagy.

Here, we show that Rab7 activity is increased in the absence of Ga_q_, consistent with a reduction in untethering events and enhanced lysosomal clustering. The observation that Ga_q_ does not directly affect the RabGAP activity of TBC1D15 suggests that its role is primarily structural or regulatory, acting as a scaffold that modulates the assembly, stability, or localization of protein complexes. Interestingly, constitutively active Rab7 recruits both Ga_q_ and TBC1D15, supporting a model in which these proteins cooperate to restrain Rab7 activity. One potential mechanism could involve the regulation of TBC1D15 protein abundance, thereby indirectly controlling Rab7-dependent processes as the present results suggest. TBC1D15 expression is known to be regulated at multiple levels, including proteasome-mediated protein turnover (Kim *et al*, 2021; Zhu *et al*, 2025) and transcriptional control by microRNA MIR1 (Ejlerskov *et al*, 2020). Consistent with this hypothesis, Yu *et al*. and Sun *et al*. reported that reduced TBC1D15 expression, either in inducible cardiac-specific knockout mice or cardiomyocytes after a myocardial infarction, leads to prolonged and enlarged mitochondria-lysosome contact sites as well as lysosomal enlargement (Sun *et al*, 2022; Yu *et al*, 2020). These phenotypes closely resemble those observed in Ga_q_-deficient cells, further supporting a functional link between Ga_q_, TBC1D15, and Rab7 in the regulation of organelle contact-site dynamics and lysosomal homeostasis.

Notably, alterations in mitochondria-lysosome contact-site dynamics have been associated with human disease and in several neurological disorders (D, PD, HD, and ALS) (Chen *et al*, 2023; Zong *et al*, 2024). In this regard, in Charcot-Marie-Tooth (CMT) neuropathy, it has been shown the increase in mitochondria-lysosome contacts and lysosomal size (Cantarero *et al*, 2023). These observations are remarkably similar to the phenotypes observed upon loss of Ga_q_ or TBC1D15 function, suggesting that dysregulation of the Ga_q_-Fis1-TBC1D15-Rab7 axis may contribute to pathological alterations in organelle homeostasis. In addition, the present findings provide a potential mechanism for the regulation of the lysosome membrane regeneration pathway, a process known to involve Rab7 and TBC1D15 (Bhattacharya *et al*, 2023). Consistent with this model, Annexin A6, a membrane-associated Ca^2+^-binding protein, promotes Rab7 inactivation through the recruitment of TBC1D15 to lysosomes and thereby influences intracellular cholesterol accumulation (Meneses-Salas *et al*, 2019). It will be important to determine whether signaling proteins such as Ga_q_ cooperate with Annexin A6 in the control of Rab7 activity and lysosomal lipid homeostasis.

The potential physiological relevance of this pathway may extend beyond organelle dynamics. Activation of a Ga_q_-coupled receptor has been proposed to reduce excessive lipolysis in adipocytes from obese mice (Kimura *et al*, 2022), thereby limiting the increase in circulating free fatty acids that promotes peripheral insulin resistance, a hallmark of type 2 diabetes. Given that enhanced mitochondrial fragmentation is a common feature of both obesity and T2D (Rovira-Llopis *et al*, 2017; Xia *et al*, 2024), the role of Ga_q_ in the regulation of mitochondria-lysosome contacts and mitochondrial fission may complement the beneficial effects of canonical Ga_q_ signaling. Thus, the combined actions of canonical and non-canonical Ga_q_ pathways may contribute to the maintenance of cellular and metabolic homeostasis, highlighting Ga_q_ as a potential therapeutic target in metabolic and other diseases.

## Methods

### Reagents

List of antibodies, cell lines, plasmids and reagents are found in Supplementary information.

### Cell culture and Transfection

Mouse Embryonic Fibroblast (MEF) cells, MEF Ga_q/11_KO (MEF knockout for Ga_q_ and Ga_11_) (a gift form Stephan Offermmans, Max-Planck-Institute for Heart and Lung Research, Germany), MEF Ga_q/11_KO stable re-expressing Ga_q_ (KI) (Benincá *et al*, 2014), MEF Ga_q/11_KO stable expressing Ga_q_E234A/E245A (KIEEAA) (produced in our laboratory from Ga_q/11_KO cells transduced cells), HEK293 cells (ATCC CRL-1573), 293-T (ATCC CRL-3216), COS7 (ATCC CCL-70), HeLa-parkin-YFP stable expressing cells (from A. Zorzano, IRB Barcelona Spain)) were maintained in DMEM GlutaMAX (BioWhittaker) supplemented with 10% fetal bovine serum (Sigma), and maintained at 37°C in a 5% CO_2_ incubator. HEK293 and COS7 were plated the day before and transfected with the desired combination of cDNA constructs using FUGENE6 (Promega) or Lipotransfectine (Niborlabs) according to the manufacturer’s guidelines. MEF cells were transfected using Lipotransfectine. Cells were tested for mycoplasma by PCR analysis or using EZ-PCR^TM^ mycoplasma detecting kit (VITRO, SA). Cell lines were authenticated by short tandem repeat (STR) or DNA barcoding analysis.

### Generation of shRNA virus and stable cell lines

pLKO.1-shRNAs to stable knock-down endogenous proteins (human Ga_q_ and mouse shFis1), and respective control vector were obtained from MISSION Sigma. Lentivirus particles were generated transfecting pLKO.1 vectors together with pCMVd-R9.91 and pVSV-G in 293-T packing cells, using Lipotransfectine (Niborlabs) according to the manufacturer’s instructions. shFis1 MEF, shFis1 Ga_q/11_KO (Ga_q/11_ KO/Fis1 KD) and shGa_q_ HeLa and HeLa-parkin stable overexpressing cells were generated. Lentivirus particles were also generated transfecting psd44-Ga_q_-E234A/E245A to generate MEF KIEEAA stable cells from MEF Ga_q/11_KO. Lentiviral purification and cell transduction were performed as described (Izquierdo-Villalba *et al*, 2024). The effectiveness of down-regulation or expression was confirmed by western blot analysis of cell extracts using specific antibodies against Ga_q_ and Fis1.

### Immunoprecipitation and western blot analysis

For western blot and immunoprecipitation, cells were lysed in buffer containing 50 mM Tris-HCl pH7.5, 150 mM NaCl, 5 mM EDTA, 1% Triton X-100, 10% glycerol, 10 mM NaF, 50 mM Na_2_H_2_P_2_O_7_, 1.5 mM MgCl_2_ with protease inhibitors as described before (Izquierdo-Villalba *et al*, 2024). GFP-Trap (Chromotek) for IP, anti-HA tag mouse Ab conjudated agarose beads (anti-HA Agarose beads, Sigma) or uncongugated antibodies together with protein G sepharose^TM^ 4 Flast Flow (GE Healthcare UK) were used for IP. Samples were resolved by SDS-PAGE before being transferred to Immobilon-FL membranes (Millipore). Primary antibodies were incubated overnight at 4 °C followed by 1 h with the corresponding anti-mouse or anti-rabbit antibodies (IRDye 680 and IRDye 800) diluted 1∶20.000 in Odyssey Blocking buffer (LI-COR). Labeling was visualized with an Odyssey infrared scanner (LI-COR). For quantification, Image Studio Software (LI-COR) was used to determine the integrated optic density (IOD). All the data was normalized by the levels of loading controls (GAPDH, Hsp90 or β-tubulin). For the proteomic analysis, data can be found in PXD046668 (ProteinXchange).

### GST pull-down assay

GST and GST-fusion proteins, GST, GST-TPR1, GST-TPR2, GST-TPR1/2, pGEX-2T-(1-200), pGEX-2T-(1-300), pGEX-2T-(304-645), were transformed into the *E. coli* strain BL21 and purified with glutathione beads (GE Healthcare). GST-proteins in 50 mM Tris-HCl pH 7.5, 150 mM NaCl, 1 mM NP-40, 0.25% deoxycholate, 1 mM EGTA pH 8.0, 1 mM NaF were incubated with lysates of HEK293 cells transiently transfected with the plasmids indicated in each figure (700 µg of protein lysates) or with MEF and Ga_q/11_KO MEF cell extracts (150 g of protein lysates) under gentle rotation for 1 h at 4 °C. Samples were rinsed with washing buffer, analyzed by SDS-PAGE, and visualized by either ODYSSEY analysis or ponceau (Sigma) staining. In pull-down experiments with purified components, 10 µg of the bead-conjugated GST, GST-Fis1 or GST-TBC1D17 proteins were incubated with 10 ng of purified Ga_q_ (Chan *et al*, 2011) in 1 ml of incubation buffer (100 mM NaCl, 20 mM HEPES, 2 mM MgCl_2_, 0.5 mM EDTA 1 mM DTT).

### Imaging of cells

Cells were seeded on 0.15-0.17-mm coverslips, fixed with 4% paraformaldehyde (15 min) at room temperature or at 37°C for mitochondria visualization, washed, permeabilized (10 min) with 0.1% triton or 0.1% saponin for lysosome visualization and blocked with 1% BSA with the corresponding detergent. Cells were incubated for 1 h with primary antibodies at the appropriate dilution in blocking buffer, then washed with PBS before being incubated with the corresponding secondary antibody at room temperature, washed in PBS, and mounted on Prolong Diamond (Invitrogen) or Mowiol (prepared according to standard protocols). Secondary antibodies (Alexa Fluor 488-, Alexa Fluor 568-, and Alexa Fluor 657-conjugated, Invitrogen) were used at a 1:2000 dilution. Mitochondria visualization was followed by staining with MitoTracker Red ROS (100nM, invitrogen), at 37°C during 30 min before fixation. Images of fixed cultures were taken on a LSM700 confocal microscope (Zeiss), epifluorescence Leica AF6500 wide-field system, spectral confocal microscope Leica STELLARIS 8 FALCON using a 63x oil immersion objective (NA 1.4) or spinning disk confocal microscope Andor Dragonfly 505 using a 60x and 100x oil immersion objectives. When needed Andor Dragonfly was used to obtain super-resolution images using the technique Super-Resolution Radial Fluctuations (SRRF) in one-z stack.

### Image processing

For the colocalization analysis in MEF WT and COS7 cells, Mander’s coefficient with threshold was calculated for each image using defined ROIs (11 µm) and the JACoP plugin from FIJI. Cell and nuclei boundaries were delineated, the nuclei were subtracted before quantification. The quantifications of the LE/Lys size, fluorescence and density in MEF WT and Ga_q/11_ KO cells were obtained by creating a mask of the LAMP1 staining and analyzing the particles size, fluorescence and number of each cell selected by ROI of the cell. Then the number of particles of each cell was divided by the area of the cell to obtain the density. The lysosome clustering in MEF WT, Ga_q/11_ KO, KI, KIEEAA and HeLa scr and Ga_q/11_ KD cells was obtained by creating a mask for LAMP1 staining, filtering the lysosomes as dispersed (0.1-2 µm^2^ and with a 0.8-1 circularity) and clustered, and calculating the percentage of the area occupied by each group respect to the total lysosomal area of the cell. The contacts between mitochondria and lysosomes in MEF WT and Ga_q/11_ KO cells were analyzed using the Manders’ Coefficients of colocalization of a square of 506.87 µm^2^. It was also analyzed in MEF WT, Ga_q/11_ KO, KI, KIEEAA and Ga_q/11_ KO/Fis1 KD cells by the creation of a mask of the LAMP-1 staining and the TOM20 staining of the same square, and counting the matching points between the two masks. The mitochondrial morphology analysis in MEF WT, Ga_q/11_ KO, KI, KIEEAA and HeLa scr and Ga_q/11_ KD cells was performed with the Mitochondria Analyzer plugging of FIJI.

### Proximity Ligation Assay

Cells were plated in 0.15-0.17-mm glass coverslips, fixed with 4% paraformaldehyde (15 min) at room temperature, washed, permeabilized (10 min) with 0.1% triton or 0.1% saponin for mitochondria and lysosomes contacts respectively, and then the Proximity Ligation Assay (PLA) utilizing the Duolink® In Situ Red Starter Kit Mouse/Rabbit (DUO92101, Sigma) was performed according to manufactures instructions. To combine the PLA with immunofluorescence after the *in situ* DNA polymerization the Alexa-Fluor 647 secondary antibody was incubated and then cells were mounted with Duolink mounting media (DUO92101, Sigma). Images were taken with a spinning disk confocal microscope Andor Dragonfly 505 using a 60x oil immersion objective. The PLA assay in MEF WT and Ga_q/11_ KO cells was analyzed by counting the number PLA puncta per cell.

### Live cell imaging

MEF WT and Ga_q/11_ KO cells (0.8×10^5^ cells/ml) plated in p35 plates (Ibidi) were incubated with 50 nM LysoTracker Deep Red (L12492, Invitrogen) and 100 nM MitoTracker Green (M7514, Invitrogen) in DMEM medium for 30 min at 37°C. Then cells were washed, and finally visualized *in vivo* with DMEM medium in a chamber with 5% CO_2_ at 37°C with the spinning disk confocal microscope Andor Dragonfly 505 immersion oil 60x objective. One-z stack was followed taking images every 2 s during 2 min. For the calculation of mitochondria-lysosome contact duration, the lysosomes trajectories were followed by the TrackMate plugin of FIJI and it was calculated the time these lysosomes colocalized with mitochondria during their trajectories.

### Mitophagy

Mitophagy was evaluated by the Mito-QC assay (McWilliams *et al*, 2016). For this, MEF and Ga_q/11_ KO cells expressing parkin and pMito-QC were incubated with 10 mM valinomicyn (Sigma) for the indicated times, fixed and proceed for confocal microscopy as described. Mitochondria were positive for GFP and mCherry whereas mitophagic particles were estimated from puncta positive only for mCherry. Quantification was done counting the % of cells that contained more than four mCherry positive puncta in a minimum of 20 cells per condition. For LC3-autophagic flux assays, cells were treated with valinomycin (10 µM), bafilomycin (200 nM, Sigma), 10-20 µM CCCP (0452, TOCRIS Bioscience), during the indicated times before lysing the cells and proceeding for western blot analysis with the anti-LC3 antibody and anti-GAPDH. After quantification with the ODYSSEY system the ratio between LC3-II to LC3-I band intensity was measured.

### Electron microscopy

Cells plated on 0.15-0.17-mm coverslips were washed with PBS and fixed with freshly made 3% glutaraldehyde solution in 0.1 M PB for 1 h at room temperature and gentle shaking. The fixation solution was renewed by fresh solution at 4 °C. Then cells were incubated with 1% OsO_4_ for 90 min at 4 °C. Samples were dehydrated in increasing ethanol solutions. Coverslips with cells were then inverted and placed on top of BEEM® capsules filled with Spurr resin. The polymerization of the resin was carried out at a temperature of 60 °C in a stove for a period of three days. At this point, the glass was removed by thermal contrast switch while the cell monolayer remained at the top of the polymerized block. The block was sectioned with a Leica ultramicrotome EM UC7, obtaining sections of 60-70 nm that were mounted on cooper grids and stained with 2% Uranyl-less solution for 10 min and with a lead-staining solution for 5 min (Parra-Ruiz *et al*, 2026). The sections were observed using a JEOL JEM-1010 transmission electron microscope coupled with an Orius SC1000 CCD camera at the Electron Microscopy Unit from the Scientific and Technological Centers of the University of Barcelona. The distance between mitochondria and lysosomes in MEF WT and Ga_q/11_ KO cells was calculated by drawing a straight line between the two organelles in the electron microscopy images and measuring the length of the line utilizing the FIJI program.

### Statistical Analyses

GraphPad Prism 10 (GraphPad) were used to analyze the data. Unpaired Student’s t test (with Welch correction when appropriate) or Mann-Whitney U test were used to test differences between two conditions. Comparison of multiple conditions was performed by one-way ANOVA with Tukey post-hoc test for parametric data or by Kruskal-Wallis test followed by Dunn’s multiple comparison test for non-parametric data. Statistical significance was set at *P* < 0.05, and represented as \**P* < 0.05, \*\**P* < 0.01, \*\*\**P* < 0.001, \*\*\*\**P* < 0.0001. Values are given as means ± SEM or in box plots indicating the median, quartiles and minimum and maximum values.

## Acknowledgements

This work was funded by the Ministerio de Ciencia e Innovación (MINECO) of Spain grants BFU2017-83379-R and PID2023-149995NB-100 to AMA, CSIC 2020AEP117 to AMA and AGAUR GRC 00421; IIV was supported by a FI fellowship from AGAUR; M.F. is supported by a FPU fellowship from MINECO, and part of the PhD Biomedicine program of UB.

## Authors contribution

A.M.A, M.F., G.G., designed and performed experiments, prepared figures, analyzed the data, wrote and revised the manuscript; M.A.C, A.A., J.R., A. B., I.I-V, performed experiments, analyzed the data, and prepared figures. M. F designed, programmed the lysosomal distribution quantification imaging assay; C. E., C.R. contributed to the acquisition and analysis of TEM experiments, and F.M. Jr. helped with study concepts and revised the manuscript, C. B. provided study concept and design, interpreted experiments, supervised the study, A.M.A. provided study concept and design, interpreted experiments, supervised the study, wrote and revised the manuscript, and obtained funding.

## Code availability

An original Fiji macro deposited in a Github repository (10.5281/zenodo.4815136).

## Contribution information

Anna M. Aragay Combas

Cristiane Beninca

