## Supplemental Figures for "Ga_q_-mediated contact release is required for efficient mitochondrial quality control"

**SUPPLEMENTARY INFORMATION**

**SUPPLEMENTARY INFORMATION:**

**Figure S1: G$\alpha_q$ associates with Fis1, TBC1D15 and TBC1D17**

**Figure S2: G$\alpha_q$ localizes with TBC1D15 and Fis1**

**Figure S3**: **G$\alpha_q$ regulates lysosome size and clustering**

**Figure S4: Increased contacts between LE-Lys and mitochondria in absence of G$\alpha_q$**

**Figure S5: Time-lapse analysis of G$\alpha_q$-dependent regulation of lysosome-mitochondria contacts**

**Figure S6: TBC1D15 co-localization with G$\alpha_q$ at the mitochondria tips**

**Figure S7: Increased fission and mitophagy in absence of G$\alpha_q$ is dependent on Fis1**

**Reagents and tool table**

**Table 1: Primary antibodies**

**Table 2: Secondary antibosdies**

**Table 3: Cell lines**

**Table 4: Reagents**

**Table 5: Plasmids**

A

### Figure S1

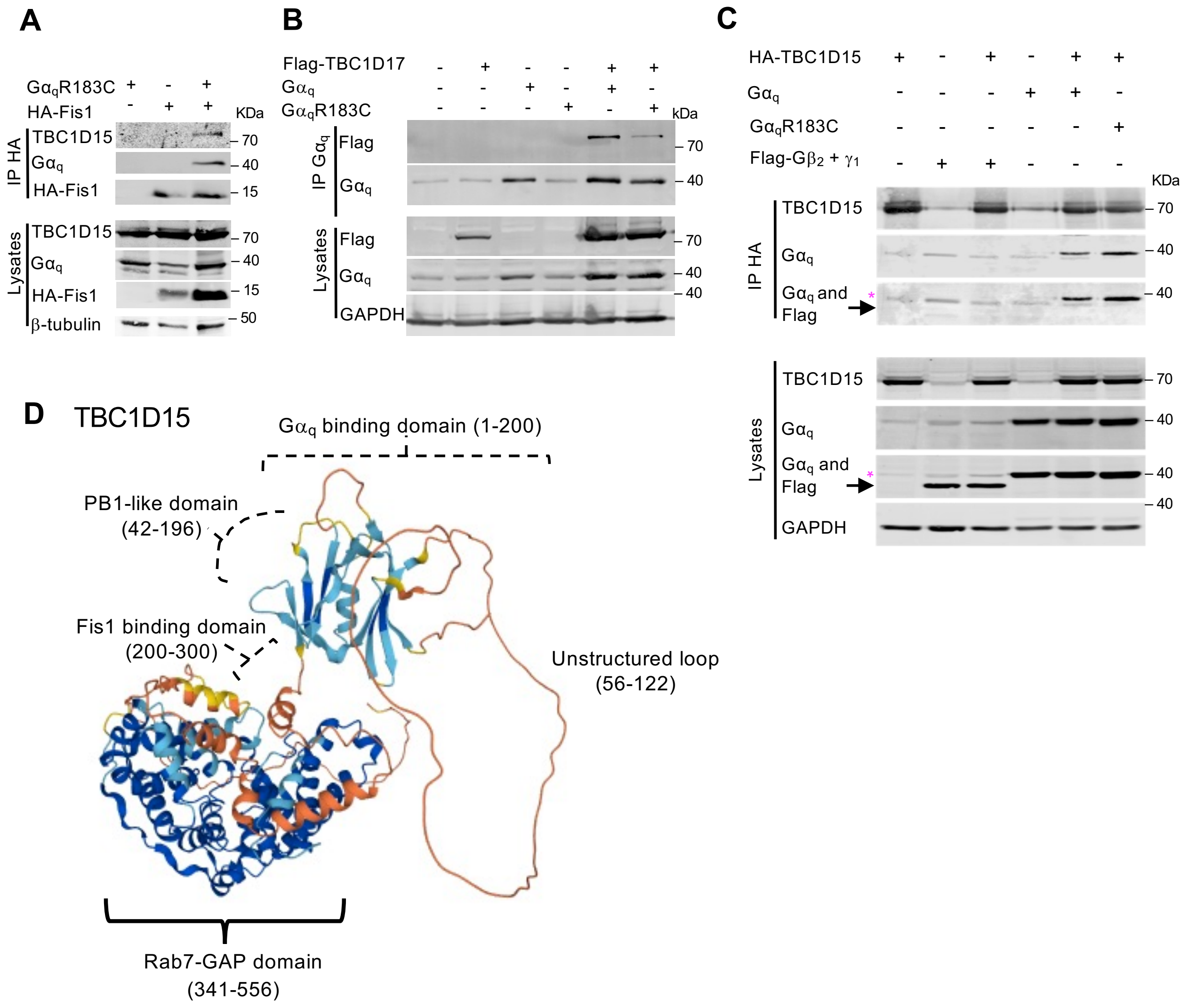
A
GαqR183C + - +
HA-Fis1 - + +
KDa
IP HA
TBC1D15 – 70
Gαq – 40
HA-Fis1 – 15
Lysates
TBC1D15 – 70
Gαq – 40
HA-Fis1 – 15
β-tubulin – 50
B
Flag-TBC1D17 - + - - + +
Gαq - - + - + -
GαqR183C - - - + - +
kDa
IP Gαq
Flag – 70
Gαq – 40
Lysates
Flag – 70
Gαq – 40
– 40
GAPDH
C
HA-TBC1D15 + - + - + +
Gαq - - - + + -
GαqR183C - - - - - +
Flag-Gβ2 + γ1 - + + - - -
KDa
IP HA
TBC1D15 – 70
Gαq – 40
Gαq and Flag – 40
Lysates
TBC1D15 – 70
Gαq – 40
Gαq and Flag – 40
– 40
GAPDH
D
TBC1D15
Gαq binding domain (1-200)
PB1-like domain (42-196)
Fis1 binding domain (200-300)
Unstructured loop (56-122)
Rab7-GAP domain (341-556)

**Figure S1. G$\alpha_q$ associates with Fis1, TBC1D15 and TBC1D17.** (**A**) HA-immunoprecipitates and lysates from HEK293 cells expressing HA-Fis1 and G$\alpha_q$R183C, immunoblotted for HA, G$\alpha_q$ and β-tubulin. Blots were also probed for endogenous TBC1D15. (**B**) G$\alpha_q$-immunoprecipitates and lysates from HEK293 cells expressing Flag-TBC1D17 and G$\alpha_q$ or G$\alpha_q$R183C, immunoblotted for Flag, G$\alpha_q$ and GAPDH. (**C**) HA-immunoprecipitates and lysates from HEK293 cells expressing HA-TBC1D15 and Flag-G$\beta_2$ and G$\gamma_1$ with G$\alpha_q$ or G$\alpha_q$R183C, immunoblotted for TBC1D15, G$\alpha_q$, Flag and GAPDH. The magenta asterisk in the IP bands indicates a nonspecific band. (**D**) Schematic model of TBC1D15 showing the G$\alpha_q$-interacting region (aa 1–200), with an unstructured loop region (aa 56-122) and PB1-like domain (aa 42-196); the Fis1-binding region (aa 200–300), and the Rab7 GAP domain (aa 341-556), based on AlphaFold predictions. All IP experiments were repeated at least three times with similar results.

### Figure S2

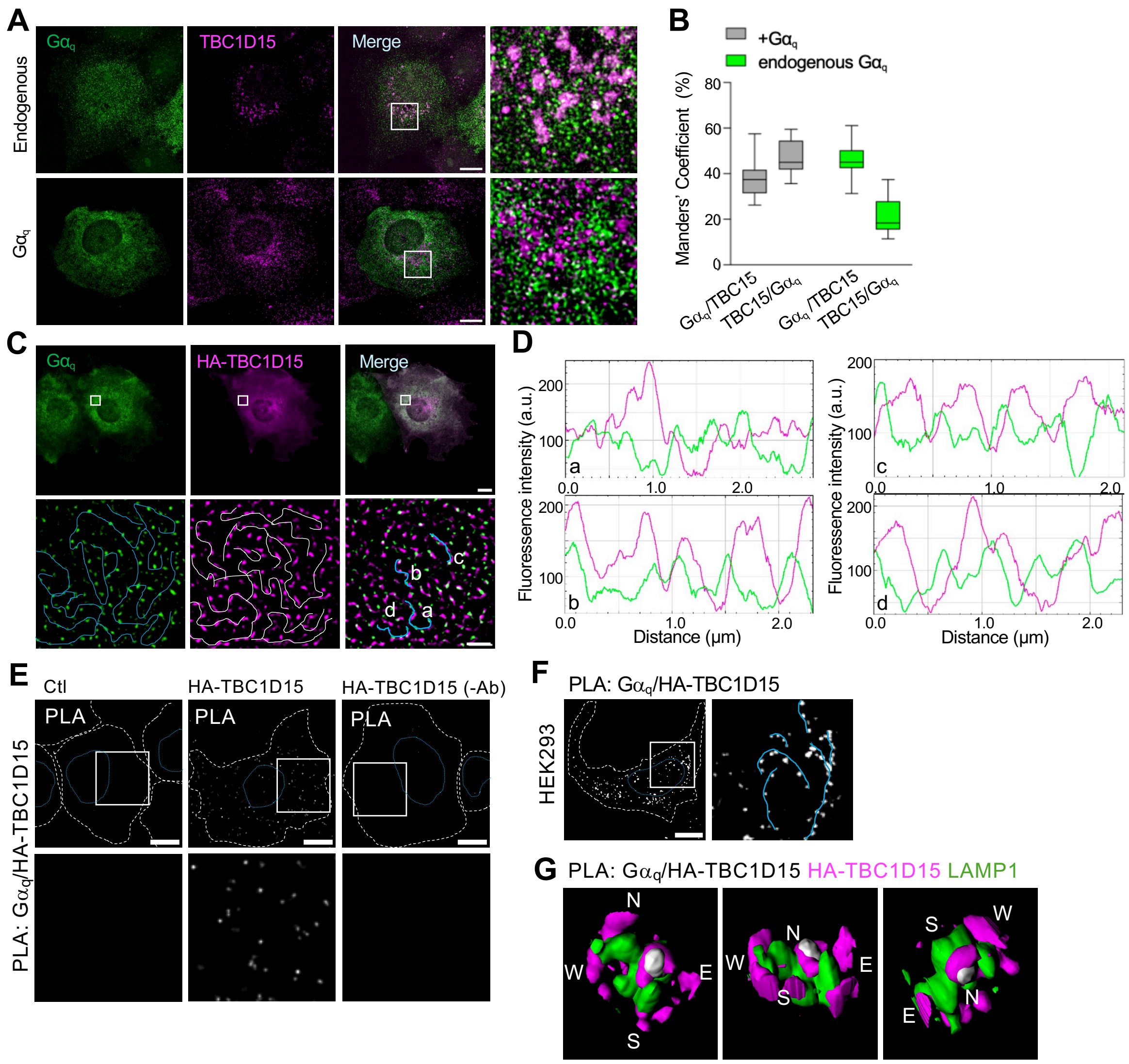
A
Gαq
TBC1D15
Merge
Endogenous
Gαq
B
+Gαq
endogenous Gαq
Manders' Coefficient (%)
Gαq/TBC15
TBC15/Gαq
Gαq/TBC15
TBC15/Gαq
C
Gαq
HA-TBC1D15
Merge
D
Fluoressence intensity (a.u.)
Distance (μm)
E
Ctl
HA-TBC1D15
HA-TBC1D15 (-Ab)
PLA
PLA: Gαq/HA-TBC1D15
F
PLA: Gαq/HA-TBC1D15
HEK293
G
PLA: Gαq/HA-TBC1D15 HA-TBC1D15 LAMP1

**Figure S2. G$\alpha_q$ localizes with TBC1D15 and Fis1.** (**A**) Representative confocal micrographs of COS7 cells untransfected (endogenous) or expressing G$\alpha_q$, immunostained with antibodies against TBC1D15 and G$\alpha_q$. Enlarged images are shown. (**B**) Manders' coefficients from images shown in (A), co-localization between G$\alpha_q$ and TBC1D15 (G$\alpha_q$/TBC15 and TBC15/G$\alpha_q$) in endogenous and G$\alpha_q$-transfected cells. $n$ = 22 cells per condition analyzed from three independent experiments. (**C**) Representative confocal micrographs of COS7 cells expressing G$\alpha_{q/11}$ and HA-TBC1D15, immunostained for HA and G$\alpha_q$. Insets show SRRF images; track marks connect individual puncta. (**D**) Fluorescence intensity line profiles along G$\alpha_q$- and TBC1D15-positive structures from images in (C). Data are representative of most cells analyzed in duplicate samples from three independent experiments. (**E**) Representative confocal micrographs of COS7 cells expressing HA-TBC1D15 and subjected to PLA using anti-G$\alpha_q$ and anti-HA antibodies. Cells without primary antibodies (HA-TBC1D15 (-Ab)) and pcDNA3 transfected cells (Ctl) served as controls. (**F**) Representative confocal micrographs of HEK293 cells expressing HA-TBC1D15 and subjected to PLA using anti-G$\alpha_q$ and anti-HA antibodies (white puncta). Track marks connect individual puncta. (**G**) 3D reconstructions of HA-TBC1D15 (magenta), LAMP1-GFP and PLA between G$\alpha_q$ and HA-TBC1D15 (white) as in (Fig. 2H) with different orientations. All scale bars are 10 µm and inset scale bars are 1 µm. All images were acquired using an Andor Dragonfly microscope and deconvolved. All images are done in duplicate samples and are representative of three independent experiments. All images the dashed white lines indicate the cell perimeter and the dashed blue the nucleus.

### Figure S3

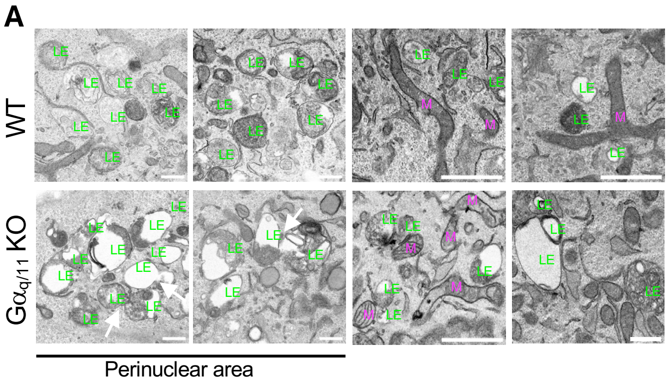


**Figure S3**. **G$\alpha_q$ regulates lysosome size and clustering.** (**A**) Four representative transmission electron microscopy (TEM) images of endolysosomes/lysosomes (LE) in WT and G$\alpha_{q/11}$ KO MEFs from perinuclear areas and other areas of the cell. Mitochondria (M) are highlated. Images taken using a JEOL JEM-1010 transmission electron microscope. Scale bars, 0.5 μm.

### Figure S4

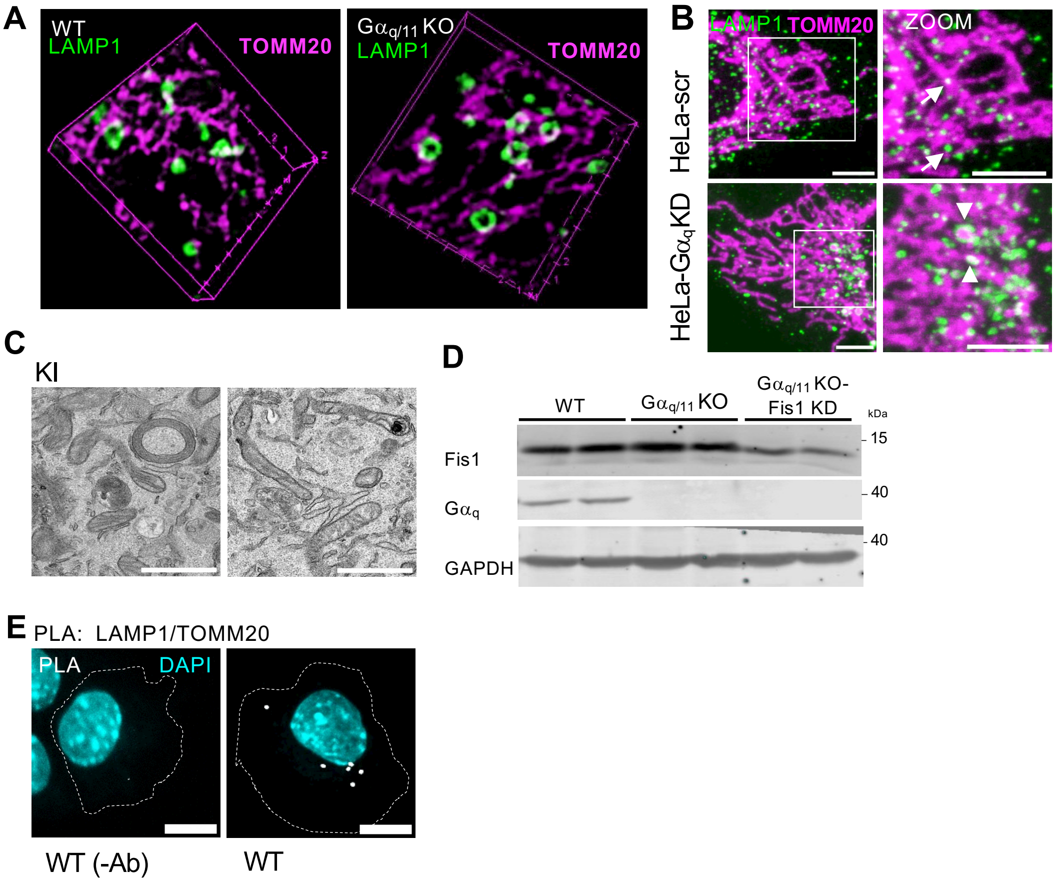


**Figure S4. Increased contacts between LE-Lys and mitochondria in absence of Gα$_q$.** (**A**) Representative confocal 3D micrographs of WT and Gα$_{q/11}$ KO MEFs immunostained for LAMP1 and TOMM20. Images were acquired using a Andor Dragonfly microscope. **(B)** Representative confocal micrographs of scramble (scr) and Gα$_q$-KD HeLa cells immunostained for LAMP1 and TOMM20. Scale bar 5 µm. (**C**) Representative TEM images showing LE–Lys (LE) in close proximity to mitochondria (M) in KI MEFs. Images were acquired using a JEOL JEM-1010 transmission electron microscope. Scale bar: 0.5 µm. (**D**) Immunoblot analysis of Fis1 expression in WT and Gα$_{q/11}$ KO MEF scramble (scr) and shFis1. Membranes were probed for Fis1, Gα$_q$ and GAPDH. (**E**) Confocal micrographs of WT MEF cells showing control PLA with (WT) and without (WT (-Ab)) antibodies and DAPI staining (cyan). Scale bar, 10 µm. Images were acquired using Leica Stellaris 8 Falcon confocal microscope (B and E).

### Figure S5

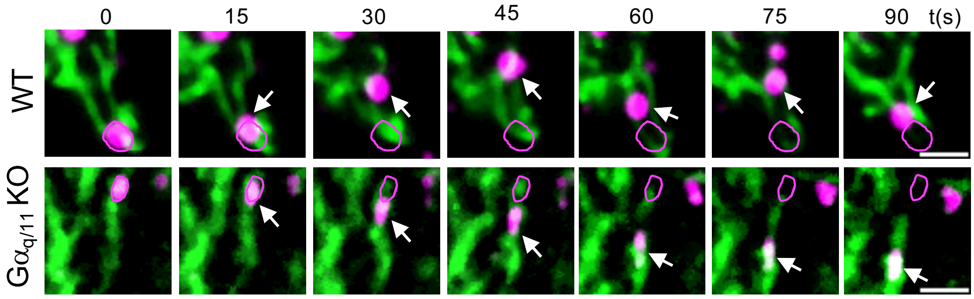


**Figure S5. Time-lapse analysis of Gα$_q$-dependent regulation of lysosome-mitochondria contacts.** Representative confocal micrographs of WT and Gα$_{q/11}$ KO cells stained with LysoTracker (magenta) and MitoTracker (green). Magenta circles mark the initial lysosome position (0 s), and white arrows indicate its position at each time point. Scale bar, 2 µm. Images were acquired using an Andor Dragonfly microscope.

### Figure S6

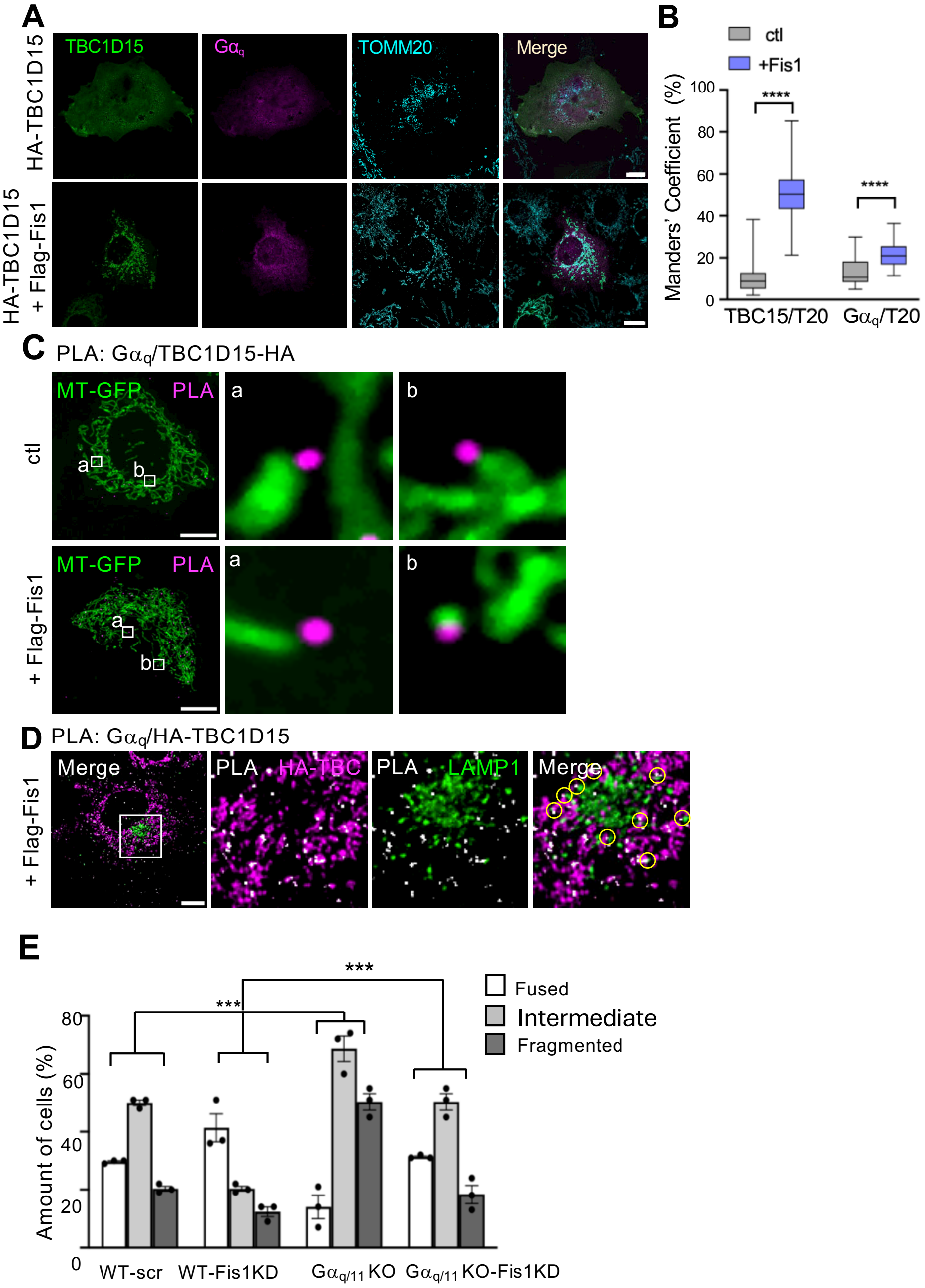
A
TBC1D15
Gαq
TOMM20
Merge
HA-TBC1D15
HA-TBC1D15 + Flag-Fis1
B
ctl
+Fis1
Manders' Coefficient (%)
100
80
60
40
20
0
****
****
TBC15/T20
Gαq/T20
C
PLA: Gαq/TBC1D15-HA
MT-GFP
PLA
a
b
ctl
+ Flag-Fis1
D
PLA: Gαq/HA-TBC1D15
Merge
PLA
HA-TBC
LAMP1
+ Flag-Fis1
E
Fused
Intermediate
Fragmented
***
***
Amount of cells (%)
80
60
40
20
0
WT-scr
WT-Fis1KD
Gαq/11 KO
Gαq/11 KO-Fis1KD

**Figure S6**. **TBC1D15 co-localization with G$\alpha_q$ at the mitochondria tips. (A)** Representative confocal micrographs of COS7 cells expressing G$\alpha_q$ and HA-TBC1D15 with endogenous or overexpressed Fis1 (Flag-Fis1), immunostained for TBC1D15, G$\alpha_q$, and TOMM20. **(B)** Manders' coefficients from images in (A) showing colocalization of TBC1D15/TOMM20 (TBC15/T20) and G$\alpha_q$/TOMM20 (G$\alpha_q$ /T20). $n$ = 38 cells per condition from three independent experiments. One-way ANOVA followed by Tukey's multiple-comparison test. **(C)** Representative confocal micrographs of WT MEFs cells expressing MT-GFP (mitochondria), HA-TBC1D15 and/or Flag-Fis1 and subjected to PLA using anti-G$\alpha_q$ and anti-HA antibodies (magenta). Two enlarged images are shown. Scale bar, 10 μm. **(D)** Representative confocal micrographs of COS7 cells expressing Flag-Fis1, HA-TBC1D15 and LAMP1-GFP (LAMP1) and subjected to PLA using anti-G$\alpha_q$ and anti-HA antibodies. PLA puncta are shown in white and HA staining in magenta. Enlarged regions are shown. Circles indicate PLA puncta associated with mitochondrial-enriched-TBC1D15- and LAMP1-positive structures. **(E)** Quantification of mitochondria morphology (%) from WT scramble (WT scr), WT treated with shFis1 (WT-Fis KD), G$\alpha_{q/11}$ KO scramble (G$\alpha_{q/11}$ KO scr) and G$\alpha_{q/11}$ KO with shFis1 (G$\alpha_{q/11}$ KO-Fis KD) MEFs stained with MitoTracker Red CMXRos. % of cells scored for completely fused, intermediate state and completely fragmented mitochondria. A minimum of 25 cells were scored for experiment and condition. Data were obtained from n = 3 independent experiments and are presented as mean ± SEM. Statistics: One-way ANOVA with Tukey. *** $P < 0.001$, **** $P < 0.0001$.

### Figure S7

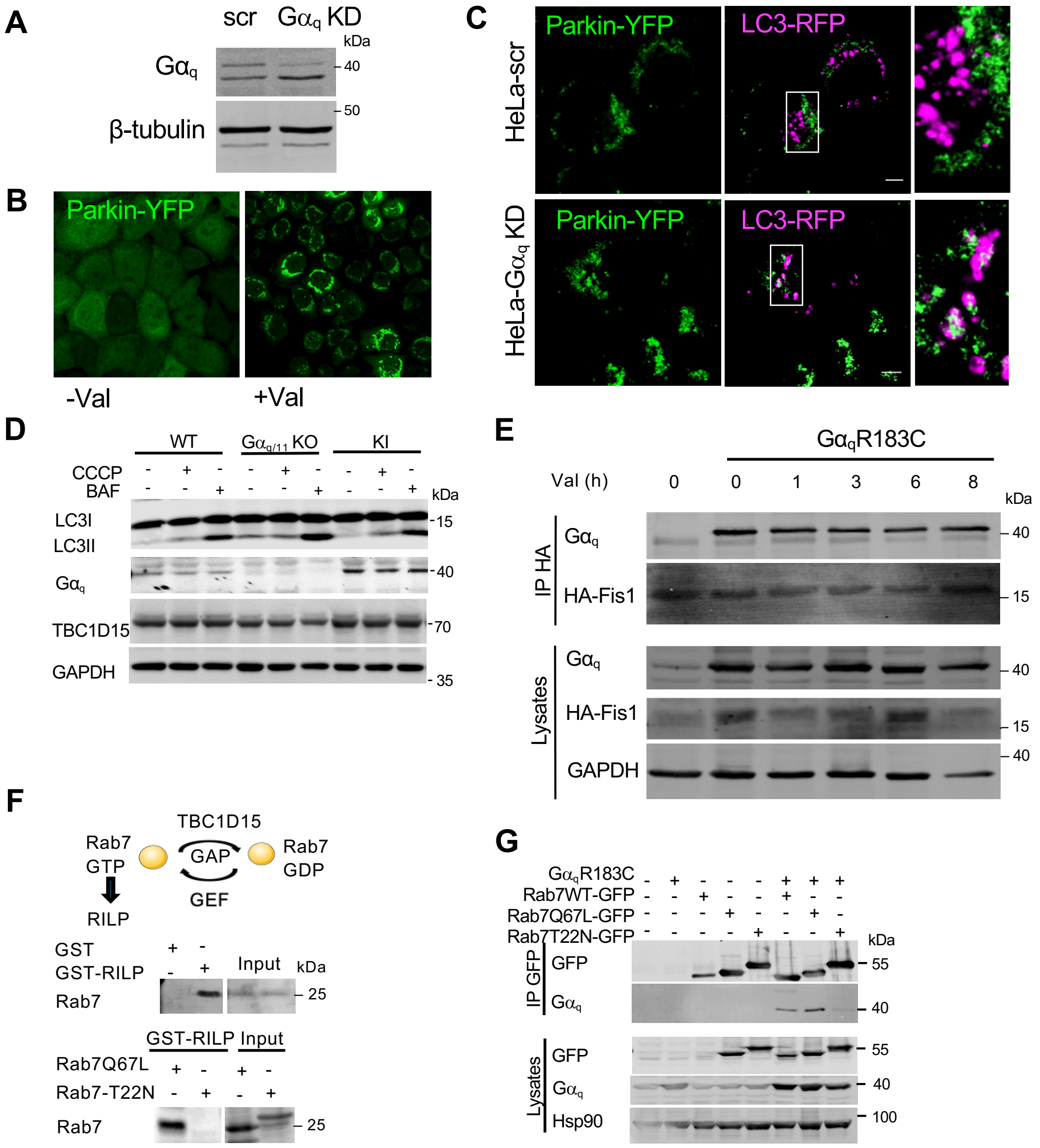
A
scr Gαq KD
kDa
Gαq
40
β-tubulin
50
B
Parkin-YFP
-Val
+Val
C
Parkin-YFP
LC3-RFP
HeLa-scr
HeLa-Gαq KD
D
WT
Gαq/11 KO
KI
CCCP
BAF
kDa
LC3I
LC3II
15
Gαq
40
TBC1D15
70
GAPDH
35
E
GαqR183C
Val (h)
0 0 1 3 6 8
kDa
IP HA
Gαq
40
HA-Fis1
15
Lysates
Gαq
40
HA-Fis1
15
GAPDH
40
F
TBC1D15
Rab7 GTP
GAP
GEF
Rab7 GDP
RILP
GST
GST-RILP
Input
kDa
Rab7
25
GST-RILP Input
Rab7Q67L
Rab7-T22N
Rab7
25
G
GαqR183C
Rab7WT-GFP
Rab7Q67L-GFP
Rab7T22N-GFP
kDa
IP GFP
GFP
55
Gαq
40
Lysates
GFP
55
Gαq
40
Hsp90
100

**Figure S7. Increased fission and mitophagy in absence of G$\alpha_q$ is dependent on Fis1.** (**A**) Extracts from HeLa cells stable expressing parkin-YFP (Parkin-YFP) transduced with virus with scramble shRNA (scr) or shRNA against G$\alpha_q$ (G$\alpha_q$ KD) were analysed by western blot and immunoblotted for G$\alpha_q$ and β-tubulin. (**B**) Confocal images of HeLa cells stable expressing parkin-YFP (Parkin-YFP) transduced with virus with shRNA against G$\alpha_q$ (G$\alpha_q$ KD) untreated (DMSO) or treated with valomycin (10 mM) for 3 h. Samples are done by duplicated in *n = 2* independent experiments. (**C**) Confocal images of HeLa cells stable expressing parkin-YFP (Parkin-YFP) transfected with RFP-LC3 and transduced with virus with shRNA against G$\alpha_q$ (G$\alpha_q$ KD) treated with valomycin (10 mM) for 3 h. Samples are done by duplicated in n=2 independent experiments. (**D**) Representative immunoblots from WT, G$\alpha_{q/11}$ KO and KI MEFs treated with CCCP (10 mM) for 3 h or with bafilomycin (BAF) (100 nM) for 5 h. Samples were immunoblotted with antibodies against G$\alpha_q$, LC3, GAPDH and TBC1D15. *n = 3* independent exprериments. (**E**) HA-immunoprecipitates and lysates from HeLa-parkin-YFP cells expressing HA-Fis1 and control vector or G$\alpha_q$R183C treated with valinomycin for the times indicated and immunoblotted for G$\alpha_q$, HA and GAPDH. *n = 3* independent expreriments. (**F**) Diagram of the Rab7 cycle illustrating the interaction of RILP with the active form of Rab7. GST and GST-RILP pull-down (PD) assay from HEK293 lysates immunostained against Rab7 (middle panel). GST-pull-down assays from HEK293 lysates expressing the Rab7-active form (Rab7Q67L) and the dominant negative for (Rab7T22N), immunostained to Rab7. Input shown on the right. (**G**) GFP-immunoprecipitates and lysates from HEK293 cells expressing GFP-Rab7, GFP-Rab7Q67L, GFP-Rab7T22N and G$\alpha_q$R183C immunoblotted for GFP, G$\alpha_q$ and Hsp90. *n = 3* independent expreriments. Images were acquired using a LSM700 confocal microscope (Zeiss).

### Table S1: Primary antibodies

| Antibody | Specie | Supplier | Catalog No. | Dilution and use |
|---|---|---|---|---|
| DsRed | rabbit | Clontech/Takara | 632496 | WB 1:2000 |
| Fis1 | rabbit | Sigma | HPA017430 | WB 1:1000 |
| FLAG | mouse | Sigma | F1804 | WB 1:1000 |
| | | | | IP 1:200 |
| GAPDH | rabbit | Cell Signaling | 14C10 | WB: 1:1000 |
| GFP | rabbit | Invitrogen | A11122 | WB 1:2000 |
| GST | mouse | Sigma | SAB4200237 | WB 1:2000 |
| $G\alpha_q$ | mouse | BD biosciences | 612704 | WB 1:1000 |
| | | | | IF 1:200 |
| | rabbit | Proteintech | 13927-1-A | WB 1:1000. |
| | | | | PLA 1:500 |
| HA (12CA5) | mouse | Invitrogen | 32-6700 | IP 1:500 |
| HA high affinity | rat | Roche | 11867423001 | WB 1:2000 |
| | | | | IF 1:500 |
| HSP-90 | mouse | BD | 610419 | WB 1:2000 |
| LAMP1 | mouse | Sigma | MABC1108 | PLA 1:500 |
| | rat | Santa Cruz | sc-19992 | IF 1:200 |
| LC3 | rabbit | MBL (Life Science) | PM036 | WB 1:1000 |
| Rab7 | rabbit | Selleckhem | F0357 | WB 1:1000 |
| TOMM20 | mouse | Sigma | WH0009804M1 | IF 1:300 |
| | mouse | Santa Cruz | sc-17764 | WB 1:1000 |
| | rabbit | Santa Cruz | sc-11415 | WB 1:1000 |
| | rabbit | Abcam | ab186734 | PLA 1:500 |
| TBC1D15 | rabbit | HPA013388 | Sigma | WB 1:1000 |
| | | | | IF 1:500-800 |
| β-tubulin | rabbit | Sigma-Merk | T2200 | WB 1:1000 |
| IRDye® 680RD Goat anti-Mouse IgG Secondary Antibody | anti-mouse | LI-COR | 926-68070 | WB 1:20000 |
| IRDye® 680RD Donkey anti-Rabbit IgG Secondary Antibody | anti-rabbit | Li-COR | 926-68073 | WB 1:20000 |
| IRDye® 800CW Goat anti-Mouse IgG Secondary Antibody | anti-mouse | Li-COR | 926-32210 | WB 1:20000 |
| IRDye® 800CW Goat anti-Rabbit IgG Secondary Antibody | anti-rabbit | LI-COR | 926-32211 | WB 1:20000 |
| Donkey anti-Mouse IgG (H+L) Highly Cross-Adsorbed Secondary Antibody, Alexa Fluor™ 488 - A-21202 | anti-mouse | Invitrogen | A21202 | IF 1:2000 |
| Donkey anti-Rabbit IgG (H+L) Highly Cross-Adsorbed Secondary Antibody, Alexa Fluor™ 488 - A-21206 | anti-rabbit | Invitrogen | A21206 | IF 1:2000 |

#### Table S2: Secondary antibodies

| | | | | |
|---|---|---|---|---|
| Donkey anti-Rat IgG (H+L) Highly Cross-Adsorbed Secondary Antibody, Alexa Fluor™ 488 - A-21208 | anti-rat | Invitrogen | A21208 | IF 1:2000 |
| Goat anti-Mouse IgG (H+L) Cross-Adsorbed Secondary Antibody, Alexa Fluor™ 568 - A-11004 | anti-mouse | Invitrogen | A11004 | IF 1:2000 |
| Goat anti-Mouse IgG (H+L) Highly Cross-Adsorbed Secondary Antibody, Alexa Fluor™ 647 - A-21236 | anti-mouse | Invitrogen | A21236 | IF 1:2000 |
| Goat anti-Rabbit IgG (H+L) Cross-Adsorbed Secondary Antibody, Alexa Fluor™ 647 - A-21244 | anti-rabbit | Invitrogen | A21244 | IF 1:2000 |

### Table S3: Cell lines

| Cell Line | Source | Catalog No. |
|---|---|---|
| COS-7 | Sigma-Aldrich | CB_87021302 |
| HEK-293 | Invitrogen | CRL-1573 |
| 293-T | ATCC | CRL-3216 |
| HeLa | Sigma-Aldrich | CB_93021013 |
| HeLa-Parkin-YFP | Prof. A. Zorzano | N/A |
| MEF wt | Prof. A. Zorzano | N/A |
| MEF KO $G\alpha_{q/11}$ | Prof. S. Offermanns | N/A |
| MEF KI (KO $G\alpha_{q/11}$ + $G\alpha_q$) | Prof. S. Offermanns | N/A |

### Table S4: Reagents

| Name | Supplier | Catalog no. |
|---|---|---|
| GFP trap A beads | Chromotek | gta-20 |
| Anti-HA Agarose Beads | Sigma-Aldrich | A2095 |
| Glutathione Sepharose Beads | GE Healthcare | GE17-5132-03 |
| DMEM with glucose and L-glutamine | Lonza BioWhittaker | 12604F |
| Fetal Bovine Serum (FBS) | Biowest | S181B |
| FuGENE 6 Transfection Reagent | Promega | E2691 |
| Lipotransfectine | Niborlabs | NL-002-01 |
| EZ-PCR Mycoplasma Detection Kit | VITRO S.A. | 20-700-20 |
| Protein G Sepharose 4 Fast Flow | GE Healthcare | GE17-0618-05 |
| Immobilon-FL Membrane | Millipore | IPFL00010 |
| Ponceau S Stain | Sigma-Aldrich | P3504 |
| Paraformaldehyde | Sigma-Aldrich | P-6148 |
| Triton™ X-100 | Sigma-Aldrich | T9284 |
| Saponin | Sigma-Aldrich | S4521 |
| Albumin (BSA) Fraction V (pH 7.0) (BSA) | Panreac AppliChem | A6588,0050 |
| Bovine Serum Albumin (IgG free) (BSA) | Jackson ImmunoResearch | 001-000-162 |
| ProLong Diamond antifade mountant with DAPI | ThermoFisher Scientific | P36966 |
| LysoTracker Deep Red | Invitrogen | L12492 |
| MitoTracker Green FM | Invitrogen | M7514 |
| MitoTracker Red CMXRos | Invitrogen | M7512 |
| Duolink In Situ Red Starter Kit Mouse/Rabbit | Sigma-Aldrich | DUO92101 |
| Valinomycin | Sigma-Aldrich | V0627 |
| Bafilomycin A1 | Sigma-Aldrich | B1793 |
| CCCP | Tocris Bioscience | 452 |
| Glutaraldehyde | Sigma-Aldrich | 49629 |
| Osmium tetroxide (OsO4) | Sigma-Aldrich | 201030 |
| Odyssey Blocking Buffer | LI-COR | 927-70001 |
| Spurr Resin | Sigma-Aldrich | EM0300 |
| Uranyl-less Stain Solution | Electron Microscopy Sciences | 22409 |
| Lead Stain Solution | Sigma-Aldrich | 18-0875 |
| Tris-HCl | Roche | 10708976001 |
| Sodium chloride | Merck | 1.06404 |
| EDTA | Sigma-Aldrich | ED2SS |
| Glycerol 85% | Merck | 1.04094 |
| Sodium fluoride | Sigma-Aldrich | S7920 |
| Sodium pyrophosphate | Sigma-Aldrich | P8010 |
| Magnesium chloride | Merck | 1.05833 |
| NP-40 | Fluka | 74385 |
| Sodium deoxycholate | Sigma-Aldrich | D6750 |
| EGTA | SERVA Electrophoresis | 11290 |
| HEPES | Sigma-Aldrich | H7523 |
| Dithiothreitol | ThermoScientific | R0861 |
| Ethanol | Carlo Erba | 4146072 |

### Table S5: Plasmids

| Plasmid | Source |
|---|---|
| pcDNA3-G$\alpha_q$ | Previously described in Benincá et al, 2014 |
| pcDNA3-G$\alpha_q$R183C | Previously described in Benincá et al, 2014 |
| pcDNA3-G$\alpha_q$-E234A/E245A | Previously described in Cabezudo et al, 2021 |
| pcDNA3-G$\alpha_q$R183C-E234A/E245A | Previously described in Cabezudo et al, 2021 |
| psd44-G$\alpha_q$-E234A/E245A | Clonned from pcDNA3-Gαq-E234A/E245A, C. Ribes (CBM) |
| pCMV3-HA-TBC1D15 | Sino Biological Inc. |
| pCMV3-Flag-TBC1D17 | Sino Biological Inc. |
| pGEX-2T-(1-300) | Clonned from pCMV3-Flag-TBC1D17 |
| pGEX-2T-(200-300) | Clonned from pCMV3-Flag-TBC1D17 |
| pGEX-2T-(304-645) | Clonned from pCMV3-Flag-TBC1D17 |
| pCMV-Flag-rFis1-p3X | K. Mihara (Tokyo Medical and Detal University, Tokyo) |
| pCEX2T | MI. Geli (IBMB, Barcelona) |
| pGEX2T-GST-TPR1 (32-61) | Cloned from pDONR223_h_Fis1, JF Rual and M. Vidal (Dana-Farber Cancer Institute, Boston) |
| pGEX2T-GST-TPR2 (62-103) | Cloned from pDONR223_h_Fis1, JF Rual and M. Vidal (Dana-Farber Cancer Institute, Boston) |
| pGEX2T-GST (32-103) | Cloned from pDONR223_h_Fis1, JF Rual and M. Vidal (Dana-Farber Cancer Institute, Boston) |
| pCDNA-HA-Fis1 | S. Grimm lab (ICL, London) |
| LAMP1-GFP | J. Lippincott-Schwartz (NIH, Bethesda) |
| pcDNA3.1-β2-FLAG | Missouri S&T cDNA Resource Center (www.cdna.org) |
| pcDNA3.1-γ1 | Missouri S&T cDNA Resource Center (www.cdna.org) |
| pLKO1-sh-scramble | MISSION Sigma (Functional Genomics Core Facility, IRB Barcelona) |
| pLKO1-shG$\alpha_q$ human (Gαq KD) | MISSION Sigma (Functional Genomics Core Facility, IRB Barcelona) |
| pLKO1-shFis1 mouse (Fis1 KD) | MISSION Sigma (Functional Genomics Core Facility, IRB Barcelona) |
| pEGFP-Rab7 | Clonned from Rab7-RFP, P. van der Sluijs lab (Utrecht University) |
| pEGFP-Rab7-Q67L-GFP | Clonned from Rab7-RFP, P. van der Sluijs lab (Utrecht University) |
| pEGFP-Rab7-T22N-GFP | Clonned from Rab7-RFP, P. van der Sluijs lab (Utrecht University) |
| Parkin-myc | A. Zorzano's lab (IRB, Barcelona) |
| Mito-GFP (MT-GFP) | A. Zorzano's lab (IRB, Barcelona) |
| Mito-QC | I. G. Ganley (U. Dundee, Scotland) |
| LC3-mRFP | A. Zorzano's lab (IRB, Barcelona) |
